# The Rise of West Nile Virus in Southern and Southeastern Europe: a spatial-temporal analysis investigating the combined effects of climate, land use and economic changes

**DOI:** 10.1101/2021.02.22.432294

**Authors:** Matthew J. Watts, Victor Sarto i Monteys, P. Graham Mortyn, Panagiota Kotsila

**Affiliations:** Institute of Environmental Science and Technology (ICTA), Autonomous University of Barcelona (UAB), Bellaterra, Spain; Departament d’Agricultura, Ramaderia, Pesca, Alimentació i Medi Natural, Generalitat de Catalunya, Avinguda Meridiana, Barcelona, Spain; Barcelona Laboratory for Urban Environmental Justice and Sustainability (BCNEJ), Institute of Environmental Science and Technology (ICTA), Autonomous University of Barcelona (UAB), Bellaterra, Spain; Department of Geography, Autonomous University of Barcelona (UAB), Bellaterra, Spain

**Keywords:** West-Nile-virus, climate-change, economic-crisis, mosquito, austerity, vector-borne-disease, drought

## Abstract

West Nile Virus (WNV) has recently emerged as a major public health concern in Europe, its recent expansion also coincided with some remarkable socio-economic and environmental changes, including an economic crisis and some of the warmest temperatures on record. Here we empirically investigate the drivers of this phenomenon at a European wide scale by constructing and analyzing a unique spatial-temporal data-set, that includes data on climate, land-use, the economy, and government spending on environmental related sectors. Drivers and risk factors of WNV were identified by building a conceptual framework, and relationships were tested using a Generalized Additive Model (GAM), which could capture complex non-linear relationships and also account for the spatial and temporal auto-correlation. Some of the key risk factors identified in our conceptual framework, such as a higher percentage of wetlands and arable land, climate factors (higher summer rainfall and higher summer temperatures) were positive predictors of WNV infections. Interestingly, winter temperatures of between 2 °C and 6 °C were among some of the strongest predictors of annual WNV infections, one possible explanation for this result is that successful overwintering of infected adult mosquitoes (likely *Culex pipiens*) is key to the intensity of outbreaks for a given year. Furthermore, lower surface water extent over the summer is also associated with more more intense outbreaks, suggesting that drought, which is known to induce positive changes in WNV prevalence in mosquitoes is also contributing to the upward trend in WNV cases in affected regions. Our indicators representing the economic crisis were also strong predictors of WNV infections, suggesting there is an association between austerity and cuts to key sectors, which could have benefited vector species and the virus during this crucial period. These results, taken in the context of recent winter warming due to climate change, and more frequent droughts, may offer an explanation of why the virus has become so prevalent in Europe.

## 1. Introduction

Over the past few decades, new health risks have been emerging in Europe, particularly with the recent appearance of neglected tropical diseases (NTDs) such as Chikungunya, West Nile Virus (WNV), Dengue (DENV-1) and Crimean-Congo haemorrhagic fever [1, 2, 3]. Rising temperatures are likely increasing the transmission potential of NTDs in Europe, by affecting the geographic spread, abundance, survival and feeding activity of vector species and benefiting pathogen development in infected vectors [4, 5, 6, 7, 8, 9]. This, combined with other factors such as human population growth, intensive animal rearing, global commerce, air travel, urbanization and land-use changes, is increasing the chances of novel diseases to enter and emerge in Europe [10, 11, 12, 13].

In this study, we focus on WNV, a single-stranded RNA *Flavivirus* closely related to other *Flaviviridae* pathogens such as dengue, Japanese encephalitis and yellow fever viruses [14]. Although WNV is a zoonotic pathogen, infecting mammals, particularly humans and horses, the transmission cycle is believed to be driven mainly by mosquitoes and birds [15], although some wild mammals may serve as intermediate hosts for West Nile virus [16]. Generally, WNV distribution is determined by the presence of suitable mosquito vectors and avian hosts, such as terrestrial and wetland birds. The spring migration of birds from infected regions of Sub-Saharan Africa to temperate regions of Europe is considered to be one of the main drivers of the disease in Europe [17, 18]. In Europe, the main vectors are *Culex pipiens*, *Culex modestus*, and *Coquillettidia richiardii*, although *Aedes* species can also transmit the disease [18] (see figures S1-S3 in additional information for vector distribution maps).

Although WNV infections are more common of late, sporadic West Nile virus outbreaks have occurred in humans and equines in southern and eastern European countries over the last century; the majority of which have occurred in wetland areas and densely inhabited urban areas [18].

Since 2010, WNV has been reported in 13 EU countries including Austria, Bulgaria, Croatia, Cyprus, Czechia, France, Greece, Hungary, Italy, Portugal, Romania, Slovenia and Spain. It has also been reported in and five EU candidate countries including Albania, Montenegro, Serbia, Turkey and Kosovo [19]. In 2010, major outbreaks hit Greece, Hungary, Romania, and Turkey. Since then, outbreaks have occurred annually in multiple regions, including more northerly regions that had not previously reported cases, including Germany, Slovakia, Slovenia, and the Czech Republic (see Figure 1). This culminated in another major outbreak in 2018, that affected more regions than had been recorded in previous years.

**Figure 1:**
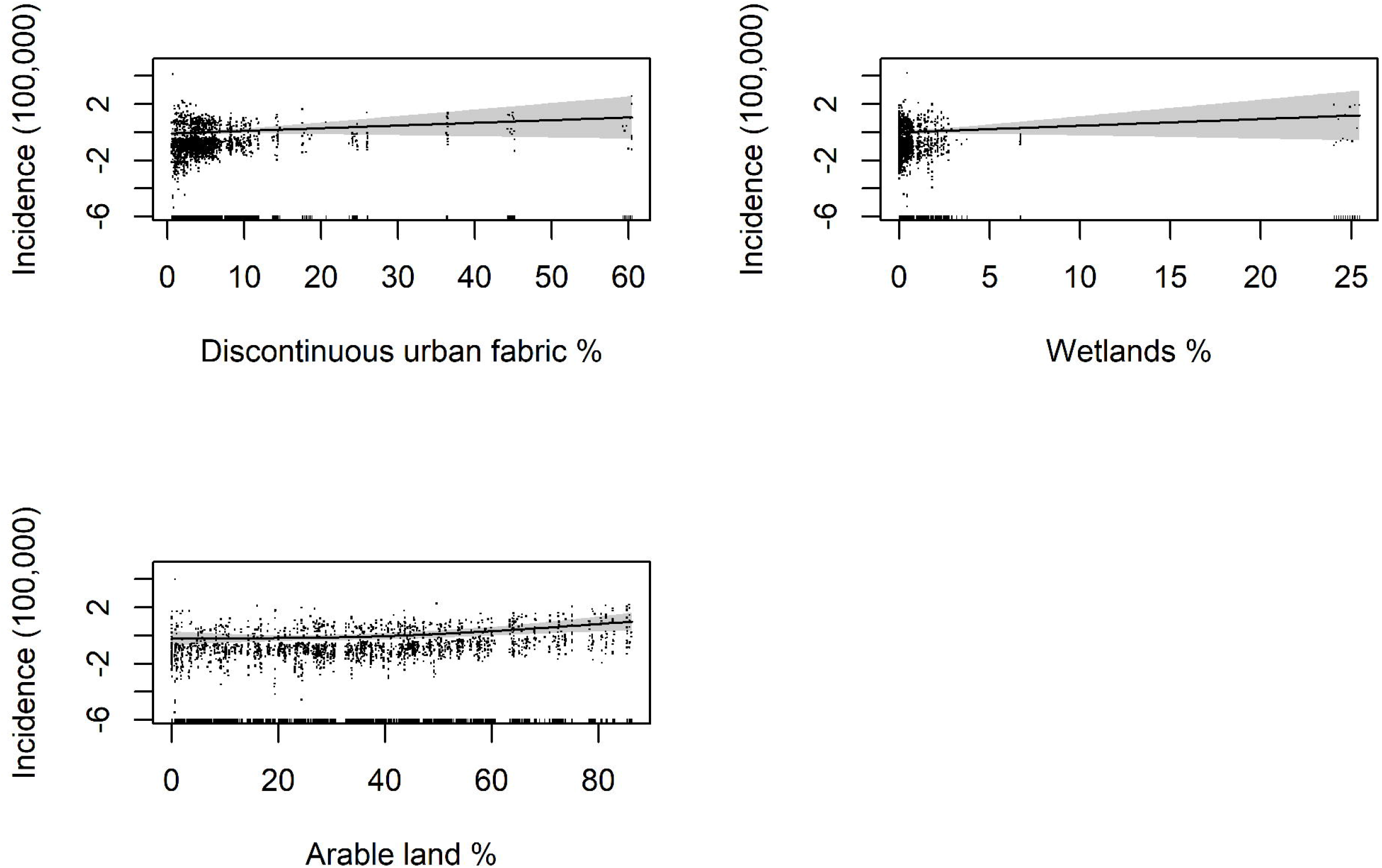
Koppen-Geiger Climate Classification in Study Regions (Data source: koeppen-geiger.vu-wien.ac.at).

We should also bear in mind that the actual number of cases in Europe is likely to be much higher than reported, since most people infected with WNV will not develop symptoms (are asymptomatic). We can therefore assume that only serious cases are reported. Around 20% of those infected with WNV will develop West Nile fever, a flu like illness, or severe West Nile disease. The most severe manifestations of the disease include acute aseptic meningitis or encephalitis, anterior myelitis, hepatosplenomegaly, hepatitis, pancreatitis, and myocarditis. On rare occasions, infection can lead to Guillain–Barré syndrome and other demyelinating neuropathies [15, 14].

### 1.1. Motivation for study

West Nile Virus (WNV) has recently emerged as a major public health concern in Europe, its recent expansion also coincided with some remarkable socioeconomic and environmental changes, including an economic crisis and some of the warmest temperatures on record. To date, there is current very little research investigating this phenomenon at a European wide scale and more work is required to reveal the key drivers of the disease. A better understanding of this phenomenon can help public health officials design health prevention measures and develop better predictive models for public health risk management. Furthermore, little work has been done to explore the association between the rise of WNV in Europe and the economic crisis. Although the study of physical factors is key in understanding disease transmission and distribution, few articles have considered how anthropogenic factors, like economic factors and policy making [20, 21], which can have wider and unintended effects on natural ecosystems and, eventually, on disease [22]. Although examining such factors presents certain challenges and uncertainties given the scales involved and lack of data. We feel there is a need to examine the statistical relationships more closely, at the very least to open up the scholarly debate and instigate further research.

To examine the recent rise of WNV infections in Europe in more depth, we empirically investigate the combined effect of three sets of factors: (1) Climate/environmental factors including temperature, rainfall and surface water; (2) land-use factors including continuous/discontinuous urban fabric, regional coverage of wetlands and arable land; (3) socio-economic factors that capture the associations of the economic crisis and are proxied by GDP growth, central government spending on areas of the environment including agriculture, forestry and fisheries and waste water management. Our analysis focuses on regions in the seven European countries where WNV has been regularly reported - Austria, Bulgaria, Croatia, Greece, Hungary Italy, and Romania. The time series data set captures the time period before and after the economic crisis (2007-2019).

Since WNV infection data is only available at the European NUTS 3 level (aggregated areal health data), our empirical strategy relies on aggregated areal health data. We recognize that the use of aggregate data presents some challenges since we cannot account for individual heterogeneity, which may lead to confounding bias i.e., relationships observed at the group level does not necessarily hold for individuals. Additionally, we may not be able to draw causal inference due to the likely presence of endogeneity bias (e.g., omitted variable bias or reverse causality). Nevertheless, we think this type of empirical investigation maintains high merit, as it allows us to quickly explore geographic associations between the disease and the predictor variables, which in turn allows us to further the debate on this topic and instigate more refined research.

### 1.2. Conceptual framework

WNV transmission requires the presence of competent vectors, a suitable climate and a susceptible host population. Studying WNV transmission at a macro scale presents significant challenges, since key data on the seasonal and annual abundance of competent mosquitoes and birds are not available, this is further complicated given the hundreds of potential host bird species in Europe [23]. To explain human infections, we therefore use environmental risk factors known to attract vector, and host. Furthermore, vector abundance for a given season is modulated by physical environmental factors, such as temperature, rainfall and water resource availability [18, 24, 14]. We therefore use proxies that can predict mosquito abundance.

#### 1.2.1. Modulating Factors

Typically, with most tropical and temperate mosquito species, elevated temperatures allow vector populations to increase their growth and reproduction rates, which in turn decreases blood meals intervals, accelerating transmission and virus evolution rates [25]. Furthermore, increasingly warmer winters allow mosquito vectors to expand their breeding seasons and survive during winter, either as eggs or as overwintering female mosquitoes. Weather conditions and climatic factors can also affect vector competence [14]. Viral replication rates and transmission of WNV are modulated by ambient temperature, affecting the length of the extrinsic incubation period (EIP), seasonal phenology of mosquito host populations and also times at which humans and mosquitoes come into contact [26]. Generally, higher rainfall in warmer weather can lead to higher mosquito abundance and disease transmission by increasing the potential habitat suitable for mosquito reproduction, e.g. standing water [27, 18, 28]. Conversely, sometimes drought and shrinking water resources can can also bring some species into closer contact, facilitating transmission and amplification of WNV within these locations [14]. To represent these points in our data-set / model, we selected mean winter temperature, mean summer temperature, number of days of rainfall in summer and summer surface water extent (the number of satellite surface water observations per region).

#### 1.2.2. Risk factors

West Nile virus circulation in Europe is usually confined to two different cycles and ecosystems: the sylvatic and the urban synanthropic cycle. Rural locations, including river deltas and floodplain areas, help create a sylvatic cycle, where wild, usually nesting wetland birds and ornithophilic mosquitoes *Culex pipiens, Culex modestus, Coquillettidia richiardii*) create the conditions for maintaining WNV transmission. In urban synanthropic cycles mosquitoes, such as *Culex pipiens* or *Culex modestus*, feed on domestic birds and humans. However, these two cycles can overlap, so areas with wetlands close to human population can be particularly at risk to the disease [18]. Irrigation from agriculture is also heavily linked to a greater incidence of human and veterinary WNV infections [29]. In order represent these factors in our data set and final models, we selected land use variables (% cover) representing urban areas (metro areas), semi urban areas (lower density human settlements), wetlands and arable land.

#### 1.2.3. Economic crisis

We would expect the repercussions of an economic crisis to affect WNV transmission in several ways, at the individual level (bottom-up) and government level (top-down). Previous studies have shown that socio-economic factors tend to influence the distribution and intensity of mosquito-borne diseases both pre-infection and post-infection [30]. Poorer communities are less likely to have air-conditioned homes, tap water and adequate drainage, and therefore may be more exposed to biting mosquitoes. Several studies have demonstrated the link between WNV infections and a range of local-level socio-economic and demographic factors such as income, sanitation, and population density [17, 31, 32]. In general, we would expect to see a drop in living standards in regions experiencing an economic shock followed by sluggish economic growth. Those people most affected would find it more difficult to prevent mosquito infections through direct measures i.e. sprays and repellents and less likely to pay for things that indirectly influence WNV transmission, like the up keep of homes and use of air conditioning. Factors associated with higher economic status can also bring humans into closer contact with mosquitoes; for example, home owners with gardens and potted plants, swimming pools and ponds or having good access to recreational space where mosquitoes can breed [33, 34]. However, neglect of such things through economic decline can have further unintended effects, even in wealthy neighborhoods [35, 36, 37].

Currently, the most effective way to prevent transmission of WNV from mosquito to human is through mosquito control (mosquito abatement). This is crucial to prevent transmission and prevent an epidemic once transmission has begun. It is well documented, for example, that during the European debt crisis, the Greek government cut mosquito abatement budgets which may have led to a rise in vector borne disease outbreaks such as malaria and WNV [38]. WNV transmission is most likely to occur in places that favor the larval development of *Culex pipiens,* such as poorly drained low-lying areas, urban stormwater catch basins and manhole chambers, roadside ditches, sewage treatment lagoons, and man-made containers around houses, or other aquatic environments where mosquitoes deposit their eggs [14]. Additionally, during periods of austerity, governments can neglect hazard prevention efforts, such as spending on flood defenses, as well as essential works like sanitation and up-keep and improvement of infrastructure [39]. Such degradation can lead to the creation of mosquito habitats [14, 18, 39]. Another critical component of preventing disease transmission is through public education programs and health promotion, educating the public on measures which can prevent being bitten can reduce risk of exposure. In general, we would expect to see a general deterioration in a government ability to run such programs during crisis and austerity. Other consequences of austerity can be expected in decreased disease detection because of cuts in public health services, prolonged periods between initial infection and treatment seeking due to dysfunctional healthcare systems, and reduced treatment of disease; all of which can lead to more intense outbreaks [40].

In order to represent the economic crisis in our model, we selected regional GDP and central government on spending on health and environment factors including agriculture, forest & fisheries and waste water management. Rather than using actual annual values, or year on year growth, we look at increases or decreases in growth using 2007 baseline levels, just before the crisis hit Europe. As a priori, we would expect WNV incidence to be associated with negative growth or very low growth in these sectors.

## 2. Materials and methods

In this study, we compiled a unique spatial temporal data-set that captures the main drivers and risk factors of WNV infections in Europe, based on findings from the conceptual framework. We selected regions for the study if autochthonous virus transmission had occurred at least once over the reporting period and if vectors were present in a region (see figures S1-3). We assumed that all regions included in the study could be influenced by migratory birds that form part of the African and European flyways [41].

### 2.1. Aggregation

All data were aggregated annually to produce the final yearly panel data-set and aggregated at the NUTS 3 country subdivision level, apart from economic data which was sourced at the country level (central government). The Nomenclature of territorial units for statistics (NUTS) is a classification system used to divide economic territories of the EU into three hierarchical sub categories for the purpose of data collection and and statistical analysis:

- NUTS 1: Major socio-economic regions with a population ranging from 3 to 7 million.
- NUTS 2: Basic regions generally used for the application of regional policies with a population ranging from 800,000 to 3 million.
- NUTS 3: Small regions for specific diagnoses with a population ranging from 150,000 to 800,000.

For further details see https://ec.europa.eu/eurostat/web/nuts/background.

#### 2.1.1. WNF Case Data

WNV case data were provided at request by the European Centre for Disease Prevention and Control (www.ecdc.europa.eu). Case data are collected weekly by EU member states and affiliates. Data were aggregated at NUTS 3 country subdivisions [42]. Positive cases were confirmed by at least one of the following techniques: 1); isolating WNV or WNV nucleic acid from blood or cerebrospinal fluid (CSF); 2) inducing a WNV-specific antibody response (either IgG / IgM) in a serological test. All cases were aggregated yearly to create the annual panel data-set.

### 2.2. Economic, Socio-Economic and Demographic

Economic data were extracted from the Eurostat database (https://ec.europa.eu/eurostat/data/database), which provides comparable statistics and indicators and is presented in yearly time series. To capture factors determining the economic crisis, austerity and cuts to public spending we selected NUTS3 regional level Gross Domestic Product (GDP); and country level agriculture, forestry, fisheries spending, waste water spending Health spending. The “Agriculture, forestry, fisheries spending” variable captures spending in rural areas that help to improve the environment and agricultural development, that can benefit agricultural workers and/or mechanize production [43]. In order to represent spending before and after the economic crisis, we created a baseline index for each variable set at 2007 levels, which represented negative or positive growth from the point just before the economic crisis hit Europe.

Population count data to predict the number of people at risk in a region were sourced from the Socio-economic Data and Applications Center’s Gridded Population of the World data set [44]. This data-set estimates population count for the years 2000, 2005, 2010, 2015, and 2020, consistent with national censuses and population registers. R’s Zoo package was used to replace values for missing years, by implementing a linear interpolation method that would predict trends between years. This way, increases or decreases in human population were controlled for in the final model.

### 2.3. Climate Data

Climate data were sourced from the E-OBS Gridded Data-set [45]. The data-set was created using a series of daily climate observations at meteorological stations throughout Europe and the Mediterranean. Files were processed in R with the Tidyverse, NetCDF and Raster packages to create regional seasonal variables: “Mean temp winter (°C)”, “Mean temp spring (°C)”, “Mean temp summer (°C)”, “Winter precipitation (total mm per region)”, “Spring precipitation (total mm per region)”, “summer precipitation (total mm per region)”. Although, to standardize the data across regions we created variables which counted rain occurrence per season i.e. “Days of rain in winter, “Days of rain in spring” and “Days of rain in summer”. Winter was designated as December to March, Spring as March to June, and summer June to September.

### 2.4. Land-use data

Land use statistics were captured at the NUTS3 level using the CORINE Land Cover (CLC) 2006, 2012 and 2018 data-sets [46]. These data-sets provide information on the biophysical characteristics of the Earth’s surface in the form of categorical raster data. For each region, we calculated percentage land cover for each of the land-use risk factors identified in our conceptual framework i.e., “Continuous urban fabric”, “Discontinuous urban fabric’, “Wetlands (fresh water)” and “Arable land”. R’s SF and Raster packages were used to extract information for each available year (2006, 2012, 2018). R’s Zoo package was used to calculate values for missing years, by implementing a linear interpolation method that would predict trends between years, apart from 2019 where 2018 values were used.

### 2.5. Surface Water data

Regional surface water data was sourced using the JRC Monthly Water History, v1.2 data set [47] via Google Earth Engine at a 30 meter pixel resolution. This data set contains maps of the location and temporal distribution of surface water from 1984 to 2019 and provides statistics on the extent and change of water surfaces. Data were generated using scenes from Landsat 5, 7, and 8. Each pixel was individually classified into water / non-water using an expert system and the results were collated into a monthly history. Surface water observations were standardized by region and season by converting them to Z-scores. This would help determine if the seasonal water extent was average, below the mean (low), or above the mean (high) for a given year.

### 2.6. Statistical Methods

#### 2.6.1. General additive regression model to assess associations of independent variables on WNV case data at regional level

One of the main issues with our data-set is that it did not meet some basic assumptions for statistical inference, and specifically the data are not independent and identically distributed random variables (iid). More specifically, the data-set captured repeated measurements over the same regions, and observations were not independent because of spill over effects from neighboring regions, therefore we needed to implement an appropriate statistical design to control for both temporal and spatial pseudo replication (lack of independence). We could deal with this in two ways, 1) either using a generalized linear mixed model (GLMM) approach, relaxing the assumption of independence and estimating the spatial/temporal correlation between residuals, or 2) model the spatial and temporal dependence in the systematic part of the model [48]. We opted to use a Generalized Additive Model (GAM) using R’s Mgcv statistical package because of its versatility and ability to fit complex models that would converge even with low numbers of observations and could capture potential complex non-linear relationships. One of the advantages of GAMs is that we do not need to determine the functional form of the relationship beforehand. In general, such models transform the mean response to an additive form so that additive components are smooth functions (e.g., splines) of the covariates, in which functions themselves are expressed as basis-function expansions. The spatial auto-correlation in the GAM model was approximated by a Markov random field (MRF) smoother, defined by the geographic areas and their neighborhood structure. We used R’s Spdep package to create a queen neighbors list (adjacency matrix) based on regions with contiguous boundaries i.e.~those sharing one or more boundary point. We used a full rank MRF, which represented roughly one coefficient for each area. The local Markov property assumes that a region is conditionally independent of all other regions unless regions share a boundary. This feature allowed us to model the correlation between geographical neighbors and smooth over contiguous spatial areas, summarizing the trend of the response variable as a function of the predictors see section 5.4.2 of [49]. In order to account for variation in the response variable over time, not attributed to the other explanatory variables in our model, we used a saturated time effect for years, where a separate effect per time point is estimated.

We first tried to fit our model using a Poisson distribution. However, the mean of our dependent variable (dengue cases by region and year) was lower than its variance - E(Y) <Var(Y), suggesting that the data are over-dispersed. We also tried to fit our models using the negative binomial, quasi-Poisson and Tweedie distribution, all particularly suited when the variance is much larger than the mean. After several tests, we concluded that the Tweedie distribution worked well with our data since it can handle excess zeros [50], and allows us to model the incident rate, although results were comparative across all distributions (note that WNV infection count data, offset by a log of population at risk was used for the neg bin and quasi-Poisson models). Analysis of model diagnostic tests did not reveal any major issues; in general residuals appeared to be randomly distributed (see additional information - figures S8-S15 and Table S1 for diagnostics).

Tweedie distributions are defined as subfamily of (reproductive) exponential dispersion models (ED), with a special mean-variance relationship. A random variable *Y* is Tweedie distributed if:

*TW*_*p*_(*μ, σ*^2^) if *Y ED*(*μ, σ*^2^), with mean = *μ* = *E*(*Y*), positive dispersion parameter *σ*^2^ and *V ar*(*Y*) = *μσ*^2^.

The empirical model can then be written as:

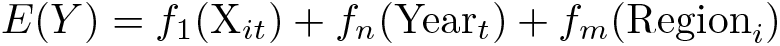

Where the *f*(.) stands for smooth functions; *E*(*Y*)_*it*_ is equal to the WNV infection incidence per 100,000 in region *i* at time *t*, which we assume to be Tweedie distributed; *Xit* - is a vector of economic, demographic, environmental and climate variables. *Y ear*_*t*_ is a function of the time intercept and *Region*_*i*_ represents neighborhood structure of region.

## 3. Results

Table 1 provides descriptive statistics of the final data-set. We did not include “Mean temp spring (°C)” in our final data set as it was correlated with winter and summer temperature variables, we concluded that we would capture more variation using the winter and summer variables which were not highly correlated. Public health spending was also not included in the final analysis as it was highly correlated with GDP (see Additional file 1 for data and model diagnostics)

**Table 1:** Final dataset - 2007-2019

| Statistic | N | Min | Max | Mean | St. Dev. |
| --- | --- | --- | --- | --- | --- |
| WNV cases | 2,158 | 0 | 100 | 1.649 | 6.012 |
| Human population | 2,158 | 6,254.326 | 10,534,640.000 | 443,384.400 | 513,000.000 |
| Mean temp winter (C) | 2,158 | -6.072 | 14.564 | 3.190 | 3.623 |
| Mean temp spring (C) | 2,158 | 4.092 | 18.684 | 12.433 | 2.157 |
| Mean temp summer (C) | 2,158 | 13.109 | 28.011 | 22.465 | 2.285 |
| Days of rain in winter | 2,158 | 0 | 68 | 30.156 | 12.546 |
| Days of rain in spring | 2,158 | 0 | 71 | 31.723 | 12.918 |
| Days of rain in summer | 2,158 | 0 | 65 | 26.021 | 14.374 |
| Spring surface water extent Z-score (30m2) | 2,158 | -2.876 | 2.404 | 0.000 | 0.958 |
| Summer surface water extent Z-score (30m2) | 2,158 | -3.301 | 3.080 | -0.000 | 0.958 |
| Continuous urban fabric % cover | 2,158 | 0.000 | 45.056 | 1.336 | 6.754 |
| Discontinuous urban_fabric (% cover) | 2,158 | 0.534 | 60.457 | 5.511 | 7.044 |
| Wetlands (% cover) | 2,158 | 0.000 | 25.460 | 0.569 | 2.026 |
| Arable land (% cover) | 2,158 | 0.000 | 86.307 | 33.893 | 22.172 |
| Regional GDP growth (2007=100%) | 2,158 | 57 | 217 | 106.752 | 24.587 |
| Agri, forest + fish spending (2007=100%) | 2,158 | 27 | 251 | 80.202 | 35.165 |
| Waste water mngmnt spending (2007=100%) | 2,158 | 5 | 352 | 93.476 | 53.933 |
| Health spending (2007=100%) | 2,158 | 60 | 212 | 114.802 | 33.307 |

To carry out our empirical analysis, we compiled a spatial-temporal dataset that captures some of the main drivers and risk factors of WNV infections in Europe. Figure 1 characterizes the climate in the study regions. As we can observe, WNV infections occurred in regions with climates that can be described as “Hot-summer Mediterranean”, “Humid subtropical”, “Temperature oceanic” and “Warm-summer humid continental” or “Temperate oceanic”. Figure 2 shows the WNV infection incidence rates over the study period. From 2007 to 2009, very few regions were affected by WNV, however, 2010 saw an outbreak that spread far and wide. Since 2010, the number of regions that have been affected by WNV increased. In 2018, a massive outbreak affected almost all of the regions in our study.

**Figure 2:**
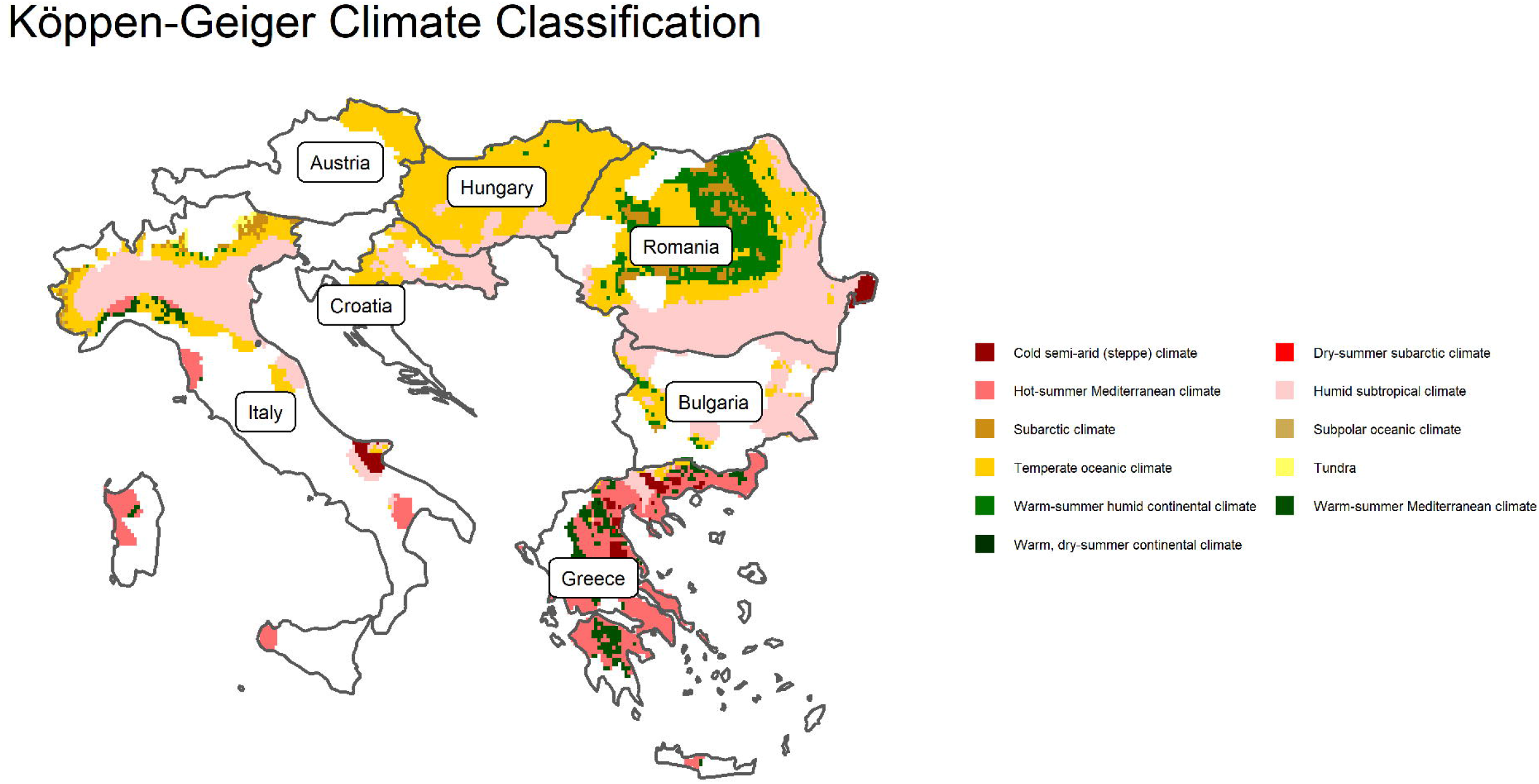
Regional West Nile Virus distribution 2006 to 2019: Confirmed cases per 100,000 (Data source: ECDC).

The relationship between the incidence of WNV infections (per 100,000) and the climate, land-use, and economic factors was modeled via a Generalized Additive Model (GAM), which also accounted for the spatial and temporal auto-correlation. Table 2 shows the results of our statistical analysis. We built the statistical model in a stepwise fashion using the lowest Akaike Information Criterion (AIC) to help us assess the different specifications. The AIC allows us to measure model performance accounting for model complexity and reflects how well the model fits the data.

**Table 2:** Final model results

|  | Clim model | Land-use model | Econ model | Full model |
| --- | --- | --- | --- | --- |
| Intercept | -2.21***<br>(0.47) | -2.13***<br>(0.45) | -2.25***<br>(0.33) | -2.35***<br>(0.40) |
| Mean temp summer (C) | 1.00***<br>(1.00) |  |  | 1.00*<br>(1.00) |
| Mean temp winter (C) | 1.96***<br>(1.99) |  |  | 1.94***<br>(1.99) |
| Days of rain in summer | 1.00**<br>(1.00) |  |  | 1.00**<br>(1.00) |
| Summer surface water extent (30m2) | 1.60**<br>(1.84) |  |  | 1.02***<br>(1.03) |
| Continuous urban fabric % |  | 1.00<br>(1.00) |  | 1.00<br>(1.00) |
| Discontinuous urban fabric % |  | 1.00<br>(1.00) |  | 1.00<br>(1.00) |
| Wetlands % |  | 1.00**<br>(1.00) |  | 1.00<br>(1.00) |
| Arable land % |  | 1.81***<br>(1.89) |  | 1.74**<br>(1.84) |
| Regional GDP index (2007=100%) |  |  | 1.00*<br>(1.00) | 1.00<br>(1.00) |
| Agri, forest + fish spending (2007=100%) |  |  | 1.95***<br>(1.99) | 1.93***<br>(1.99) |
| Waste water managment spending (2007=100%) |  |  | 1.60***<br>(1.83) | 1.10***<br>(1.19) |
| Spatial lag | 78.44***<br>(109.60) | 80.54***<br>(111.42) | 88.90***<br>(121.08) | 76.19***<br>(106.52) |
| Year | 11.73***<br>(12.00) | 11.75***<br>(12.00) | 11.48***<br>(12.00) | 11.56***<br>(12.00) |
| AIC | 3952.99 | 3992.59 | 3931.25 | 3907.56 |
| BIC | 4526.68 | 4569.26 | 4560.01 | 4538.85 |
| Log Likelihood | -1875.44 | -1894.71 | -1854.87 | -1842.57 |
| Deviance | 3747.15 | 3895.52 | 3592.25 | 3520.85 |
| Deviance explained | 0.63 | 0.62 | 0.64 | 0.65 |
| Dispersion | 2.85 | 2.95 | 2.76 | 2.73 |
| R<sup>2</sup> | 0.26 | 0.36 | 0.32 | 0.25 |
| GCV score | 1900.89 | 1927.27 | 1889.38 | 1871.90 |
| Num. obs. | 2158 | 2158 | 2158 | 2158 |
| Num. smooth terms | 6 | 6 | 5 | 13 |

\*\*\* $p < 0.01$ ; \*\* $p < 0.05$ ; \* $p < 0.1$

We selected relevant variables in each specification according to their category, i.e. climate, land-use and economic. All variables were included in the final specification to ascertain the contribution of each driver, all else equal. Table 2 also summarizes the relevant statistics (AIC, BIC and Deviance explained and so on) to compare the different specifications. We find that our final model (Full model) has the best fit in terms of the AIC, followed by the economic model, the climate model and land use model, as shown in Table 2.

Note that as we are not estimating a standard regression model, the figures reported should not be read as coefficients, but degrees of freedom of the smooth terms. Given that we cannot interpret the coefficients to infer the sign and magnitude of the relationship, we visualize it by plot. Figures (3–5) plot the partial effects—the relationship between a change in each of the covariates and a change in the fitted values in the full model.

**Figure 3:**
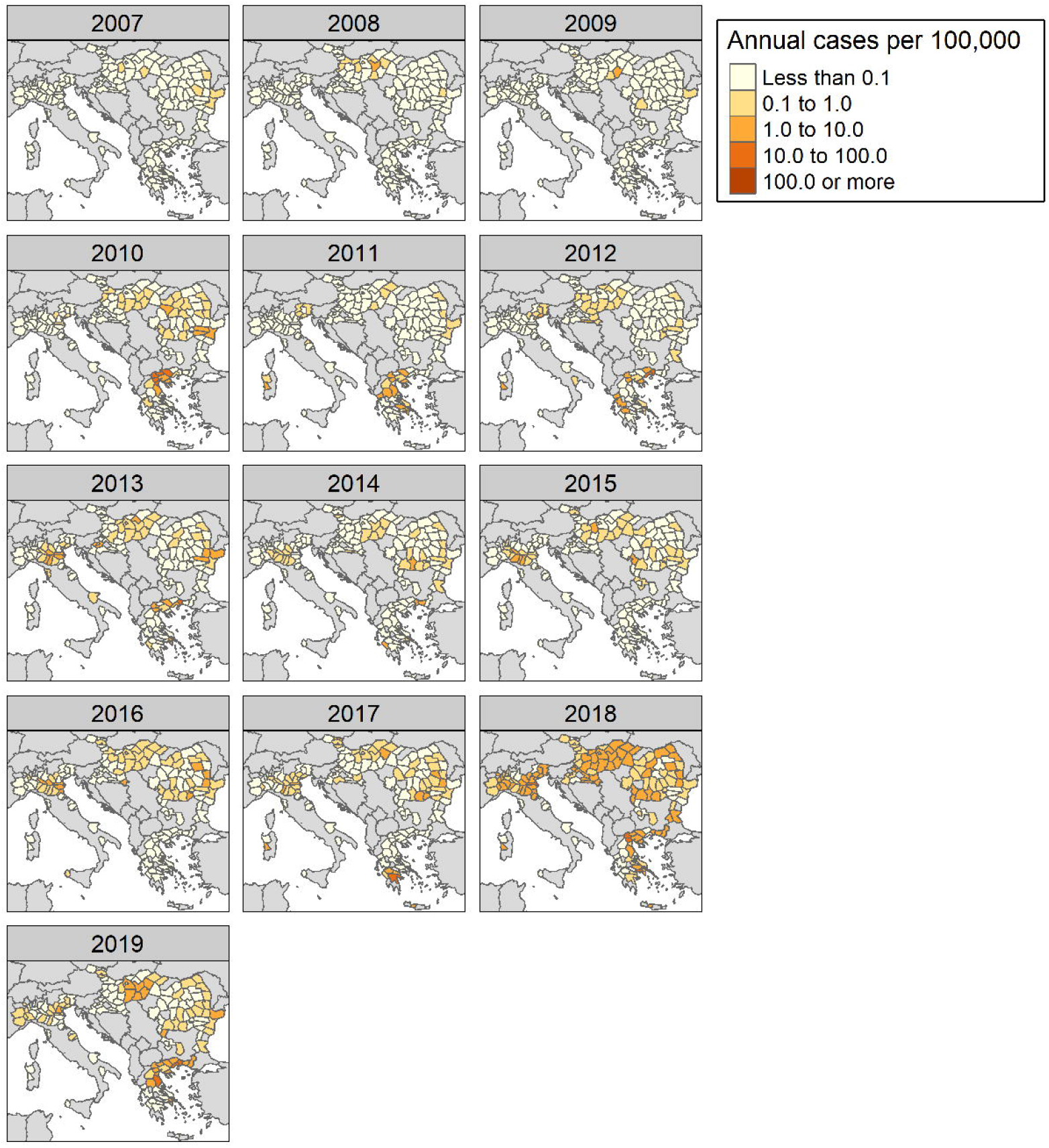
Partial effects plots of main model results 1: represents the relationship between a change in each of the covariates and a change in the fitted values in the full model.

**Figure 4:**
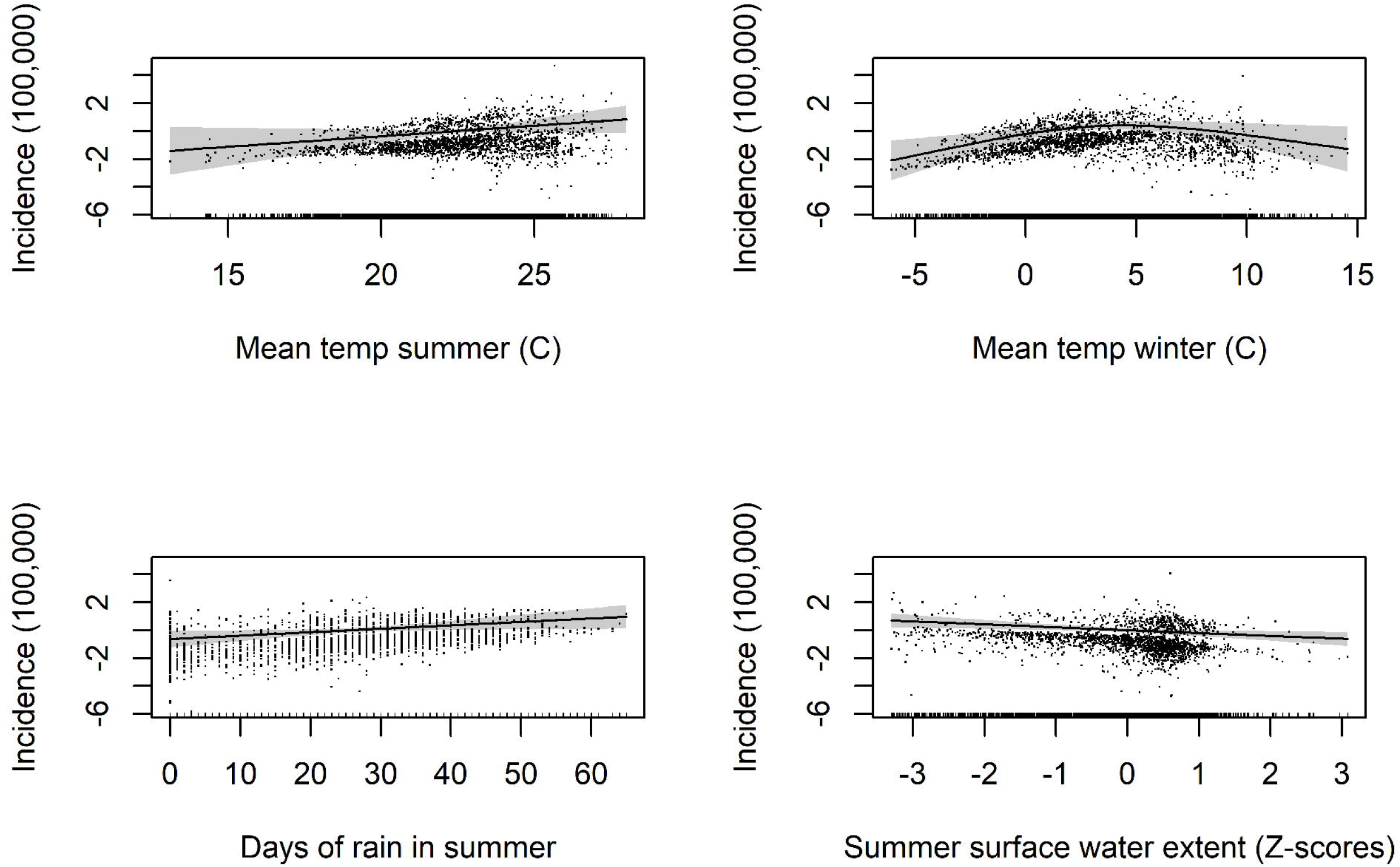
Partial effects plots of main model results 2: represents the relationship between a change in each of the covariates and a change in the fitted values in the full model.

**Figure 5:**
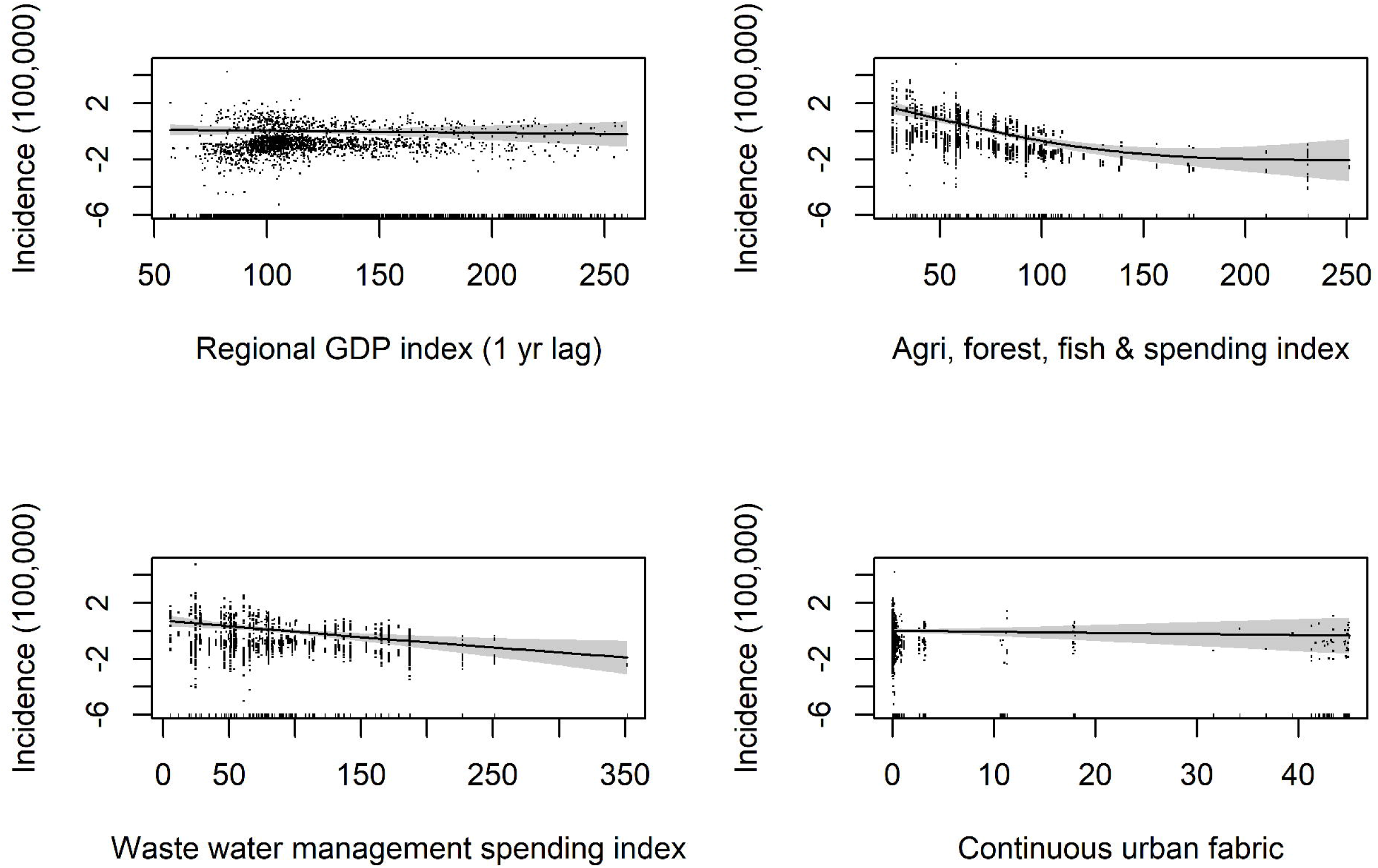
Partial effects plots of main model results 3: represents the relationship between a change in each of the covariates and a change in the fitted values in the full model.

All climate variables in both model specifications are highly statistically significant (“Clim model” & “Full model”). Mean summer temperatures of above 22°C are (Figure 3, top left) positively associated with WNV infections, the relationship is linear and strong.

Mean winter temperature has a quadratic relationship with WNV. Temperatures of between 2 °C and 6°C have a positive association with WNV infections, whereas colder and warmer temperatures outside of this range are negatively associated with WNV infections. The number of rain days per summer is also a strong predictor of WNV infections and has a linear positive relationship. Surface water in the summer is negatively correlated with WNV infections. The relationship is fairly strong considering its complexity i.e. the variable complexity has been reduced to standard deviation scores to standardize it across regions and seasons.

As for the land use variables, the percentage of arable land and wetlands in a region (“Land-use model”) is positively correlated with the incidence of WNV and highly significant. However, the wetlands variable loses significance in the final model. The percentage of “Continuous” and “Discontinuous urban fabric” variables, which represents metropolitan and built up areas (residential suburbs, villages) are not statistically significant in any of the specifications. Although the partial effect plot for “Continuous urban fabric” reveals that it has slightly negative relationship with WNV infections and “Discontinuous urban fabric” has a positive association with WNV infections.

The economic indicators are negatively correlated with WNV infections. These variables represent 2007 baseline regional GDP and central government spending growth. Regional GDP has a gentle negative association with WNV infections, but loses significance in the final model. The two indicators that directly represent central government spending on areas of the environment, such as agriculture, forest & fisheries spending and waste water management, have very strong negative associations with WNV infections.

## 4. Discussion

To help us explain the rise in WNV infections in Europe over the last 13 years (2007-2019). We compiled a unique spatial-temporal data-set that merged a range of regional abiotic factors associated with human WNV infections, along with a set of economic indicators that captured the economic crisis. This allowed for the identification of changes, in both environmental and economic factors that possibly led to the recent rise in infections over the past decade or so.

### 4.1. Meteorological factors

Over the past 70 years, the countries analyzed in this study have been experiencing increasingly warmer temperatures throughout the year (see Additional information: figures S4-7). According to our initial analysis, the last decade has been the warmest. The results of our final model (Figures 3–5) show that average summer temperatures above 22 °C are positively associated with an increase in WNV incidence. It is likely that rising temperatures have created more favorable conditions for the virus and this is why it is occurring in more northerly regions than in previous decades. Higher temperature, especially in summer months, is a key driver of WNV transmission, since warmer temperatures influence the hatching rate and development time of mosquitoes, and shorten the extrinsic incubation period (EIP) of WNV and related viruses [51, 52, 53, 54, 55, 26].

Our analysis of the mean winter temperature (Dec-Feb) reveals a quadratic relationship with WNV and is one of the strongest predictors of annual WNV infections. Temperatures below 2 °C and above 6 °C have a negative association on WNV infections. These results are consistent with findings by Koenraadt et al., 2019 [56]. The authors looked at *Culex pipiens* overwintering potential in mosquitoes in sheds and houses. Temperatures in sheds ranged from 2.7–17.0°C, and in houses 8.8–23.7°C. Survival rates were much better in sheds that had cooler temperatures. Diapausing *Culex pipiens* mosquitoes do not necessarily do better under warmer conditions and there is a temperature range in which they can successfully diapause. Although our results may be slightly less accurate than results derived through an experimental study (since we analyzed infection data at a regional scale using mean monthly temperatures), as with the study by Koenraadt et al. [56], the results do adhere to the same quadratic relationship. This can be interpreted as WNV infections for the months following winter are higher, if the specific temperature range of between 2 °C and to 6 °C is met. This is likely because WNV can overwinter with the mosquitoes when conditions are right, and then can be transmitted early on in the year, and throughout the year. This result, taken in the context of recent winter warming due to climate change, may explain why the virus has become so prevalent in Europe. Given current trends, we can also expect to see regions that have previously been too cold for *Culex pipiens* to survive over winter become viable locations and cause further havoc in regions that are currently experiencing just a few annual cases. On the contrary, regions which currently have optimal conditions for overwintering mosquitoes may become too warm. It is also important to note that it is not currently clear at what temperatures *Culex modestus* and *Coquillettidia richiardii* overwinter as adults, since our literature search didn’t yield any findings.

The number of rain days per summer is also positively correlated with WNV infections. This result is consistent with the literature, i.e., a steady flow of aquatic resources for mosquitoes has a positive association on their abundance, and therefore an increase in disease transmission [28, 27]. Rainfall patterns have also been shifting since the 1950s (see Additional information: figures S4-7) although unlike climate, there is no a clear trend and results are more difficult to interpret. In general, Austria, Croatia and Italy are seeing less intense rainfall in the summer months, but higher in autumn, whereas Bulgaria, Greece, Hungary and Romania are receiving more rainfall in summer months.

Our results also show that higher regional summer surface water extent for a given year, is negatively correlated with WNV and is one of the strongest predictors of WNV incidence. This was not an expected finding, since we would expected higher levels of surface water to be positively correlated with WNV incidence because of the extra water resources available to mosquitoes. However, the result is consistent with the literature as sometimes desiccation of water resources can bring mosquito and bird hosts closer together, increasing transmission potential and therefore the prevalence of the virus. [18, 14]. This was also a major finding in a recent study by Paull et al., 2017 [57], who reported that drought was closely linked to the intensity of outbreaks for a given year in the United States. It may be the case that this phenomenon is also acting at a macro-scale in Europe and is a significant driver of recent outbreaks, especially given that meteorological and hydrological droughts are becoming more frequent and extreme [58]. Another explanation for this result is that, with higher surface water extent may be associated with flooding and fast water movement, which may wash away mosquito eggs and larvae, and also may inhibit contact between birds, mosquitoes and humans [18, 14]. It is important to note that this variable probably does not capture the creation of short-term water resources created by rainfall (e.g. pools, puddles), which can be used as breeding habitat by mosquitoes. It rather captures long term and large water surface such as deltas, lakes and flood plains.

#### 4.1.1. Land-use

As for the land-use variables, as expected, regions with a larger proportion of arable land (including rice paddies and irrigated agriculture) and wetlands are associated with WNV incidence. This is consistent with the literature as such land-use patterns attract bird hosts and mosquito species capable of transmitting WNV, which can then have knock-on effects on the human population. The percentage of discontinuous urban fabric, that represents populated areas of low to medium density such as residential suburbs, villages etc. [59] is positively associated with WNV incidence, which is consistent with the literature, however, this variable is not statistically significant.

#### 4.1.2. Economic-factors

In terms of economic factors associated with WNV infections, lower GDP growth, lower spending growth on environmental factors such as agriculture, forest, fisheries, and waste water management are positively associated with WNV incidence. Although GDP is a very broad measure, it does act as a proxy, albeit a weak one for population health and well-being as laid out in the conceptual framework. It may be the case that populations living in locations that were harder hit were more exposed to mosquitoes for reasons outlined in the conceptual framework i.e. direct and indirect interventions i.e., drops in income make it difficult to afford mosquito repellents, air conditioning and upkeep of homes leading to the creation of mosquito habitat. Waste water management spending may be a reliable causative factor, due to, for instance, the neglect of hazard prevention efforts, such as spending on flood defenses, essential works like sanitation and upkeep of infrastructure, that can lead to the creation of mosquito breeding habitats, e.g. potholes, blocked drains [14, 18, 39]. Lack of central government spending in this area may have led to a general degradation of infrastructure and benefited mosquito species. Agriculture, forest + fisheries sectors, which represents investment or subsidies, that can go into improving environmental care, landscapes and biodiversity in rural locations, as well as investments in agricultural technology that improve aspects of the environment, worker safety or mechanize agriculture, also had a negative association with WNV infections. Many studies report strong associations between agriculture [60, 29, 61, 62] and WNV incidence, in general, cuts and lower spending in this sector may have led to degradation on farms and the wider environment which may have benefited mosquitoes through the creation of habitat or lack of measures to control their abundance. Although it is difficult to lay out actual mechanisms when dealing with data at this scale, the overall take home message here is lower annual growth in the areas i.e. economic slowdown and austerity, is positively associated with WNV incidence.

#### 4.1.3. Conclusions

In this study we set out to investigate why WNV outbreaks have become so frequent in Europe over the past decade. Our findings suggest that this phenomenon occurred because of a combination of environmental changes: 1) Rising winter temperatures, or rather the creation of optimal temperature conditions allowed the virus to overwinter with *Culex pipiens*; 2) Warmer summer temperatures are benefiting mosquitoes, influencing their hatching rate and development time, and shortening the extrinsic incubation period (EIP) of WNV; and 3) Shrinking water resources are increasing in WNV prevalence in birds and mosquitoes during some seasons. These changes also occurred during an economic crisis and subsequent austerity, where government institutions were severely weakened and had to limit spending on key sectors, and segments of the human population were exposed to increased financial hardship.

#### 4.1.4. Limitations

Some of the limitations of the study are as follows. Since we used data-driven approaches to investigate how aspects of the economy, environment and climate drive WNV incidence at a macro-scale, we were limited to using aggregated data at the NUTS-3 regional level. With any analysis dealing with population-level areal data, issues can arise due to individual heterogeneity, which may lead to confounding bias, and we are not able to draw causal inference due to the likely presence of endogeneity bias (e.g., omitted variable bias or reverse causality). Therefore, we carefully evaluated results from this study with individual-level and clinical based studies to draw conclusions. We would have also liked to include further explanatory variables on avian host and mosquito abundance but were restricted by the availability of data.

It is also important to note that data quality issues arise owing to the under-reporting of cases i.e., through under-diagnosis, lack of diagnostic tests and a lack of resources/time to carry out and implement mass testing. Another factor we did not consider is bird immunity, which may influence WNV incidence following a major outbreak, although this was not considered an important factor in explaining the rise in WNV infections in Europe, but may have influenced the results. Furthermore, a growing body of literature reports that mammals can serve as intermediate hosts for West Nile virus [16] and more research needs to be done on to determine if wild mammals act as reservoirs and contribute significantly to the transmission cycle. We also realize that the economic analysis is slightly limited, in part because of a lack of refined data and in part because scale issues i.e. the amount of work required to look at individual local level policies and spending was no feasible for 166 regions. Nevertheless, we hope this study would spur further research into this topic i.e., the impacts of the economic crisis on health; and the long term trade-offs and unintended consequences austerity can they have on the environment and human health. This is an especially important topic when considering we are facing multiple threats brought about by global warming and other anthropogenic induced changes that can benefit emerging diseases, i.e., global trade in wild animals, intensive agriculture / animal rearing and land use conversion.

## Acknowledgments

This research was funded by ICTA’s Maria de Maeztu Unit of Excellence, awarded by the Spanish Ministry of Economy and Competitiveness. The award is the highest institutional recognition of scientific research in Spain.Thanks also to Patrizia Ziveri and Pedro Manuel Gonzalez Hernandez for supporting the project. Thanks also to Cesira Urzi Brancati for statistical help and advice.

## Author contributions statement

MW led the work and was responsible for the conceptualization of the project, data collection, data processing, formulation of the methodology, statistical analysis, modeling, analysis and writing the original draft and interpreting the results. All authors made contributions to developing the original paper outline. PK gave critical advice throughout the project. All contributors revised the manuscript and copy-edited the final submission version. All the authors were also involved in revising the manuscript critically for important intellectual content. All authors read and approved the final manuscript. PK, PGM VSM supervised the project and are responsible for formulating ideas for the umbrella project Impacts of Climate Change (CC) on Human Health (HH) at ICTA-UAB: Integrating socio-economic and policy studies with natural science studies to enhance consilience of climate policy science.

## Data availability

An R project containing all the data that supports the findings of this study are available in .Rdata format from https://doi.org/10.5281/zenodo.4656902).

## Code availability

An R project containing all the code used to set-up the models is available here is available from https://doi.org/10.5281/zenodo.4656902.

