## Supplementary material for "The Rise of West Nile Virus in Southern and Southeastern Europe: a spatial-temporal analysis investigating the combined effects of climate, land use and economic changes"

### The Rise of West Nile Virus in Europe: Supplementary Information

Matthew J. Watts et al., (2021)

#### Climate modeling

In order to model long term seasonal climate trends, we fit a GAM model using the following equation.

$$y = \beta_0 + f(x_1, x_2) + \varepsilon$$

where $y =$ is either the mean of the monthly regional temperatures (°C) or regional sum precipitation (mm).

$B_0$ is the intercept, month is represented by $x_1$ and $x_2$ is the series of years in the entire time period i.e. within-year and between year.

$f$ is a smooth function interaction that accounts for variation in, or interaction between, the trend and seasonal features of the data.

Temperature models were fit using the Gaussian distribution and precipitation models fit using the Tweedie distribution.

S

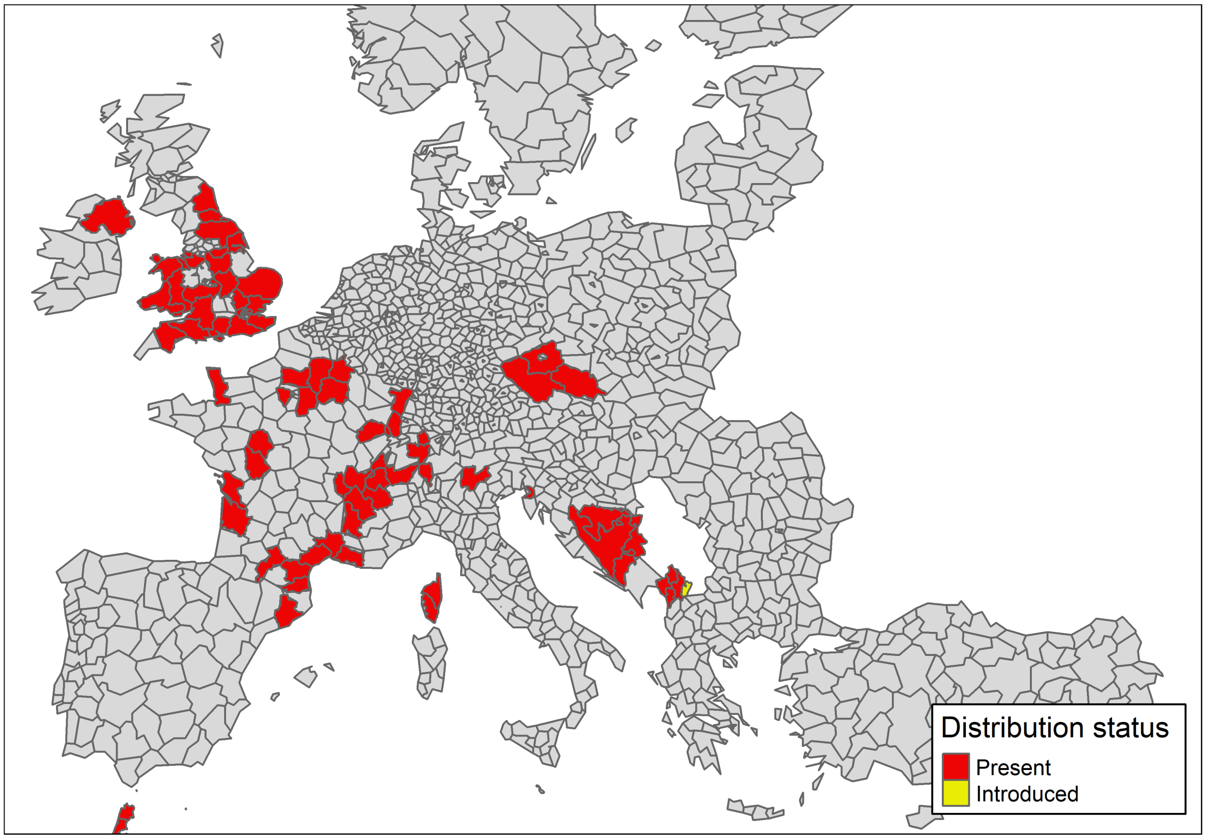


Figure S1: Confirmed Coquillettidia richiardii distribution (source: ECDC).

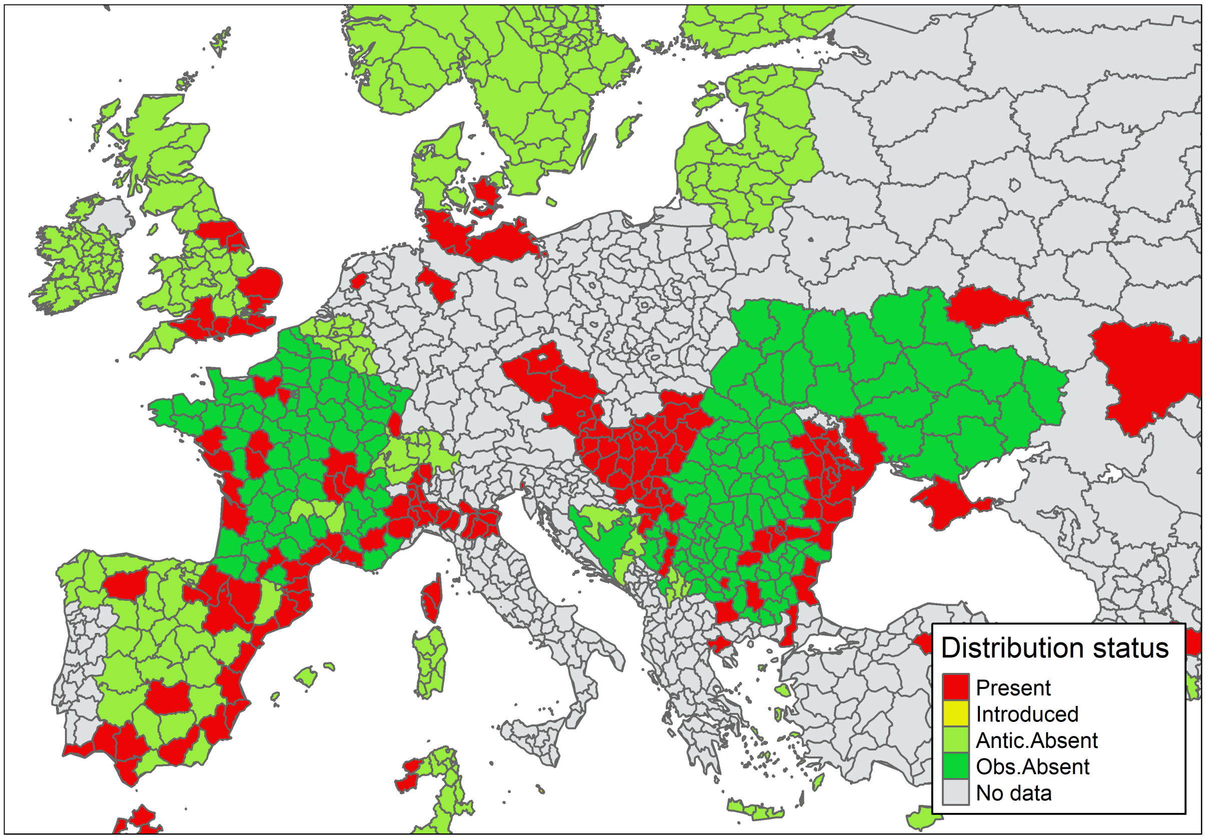


Figure S2: Confirmed Culex_modestus distribution (source: ECDC).

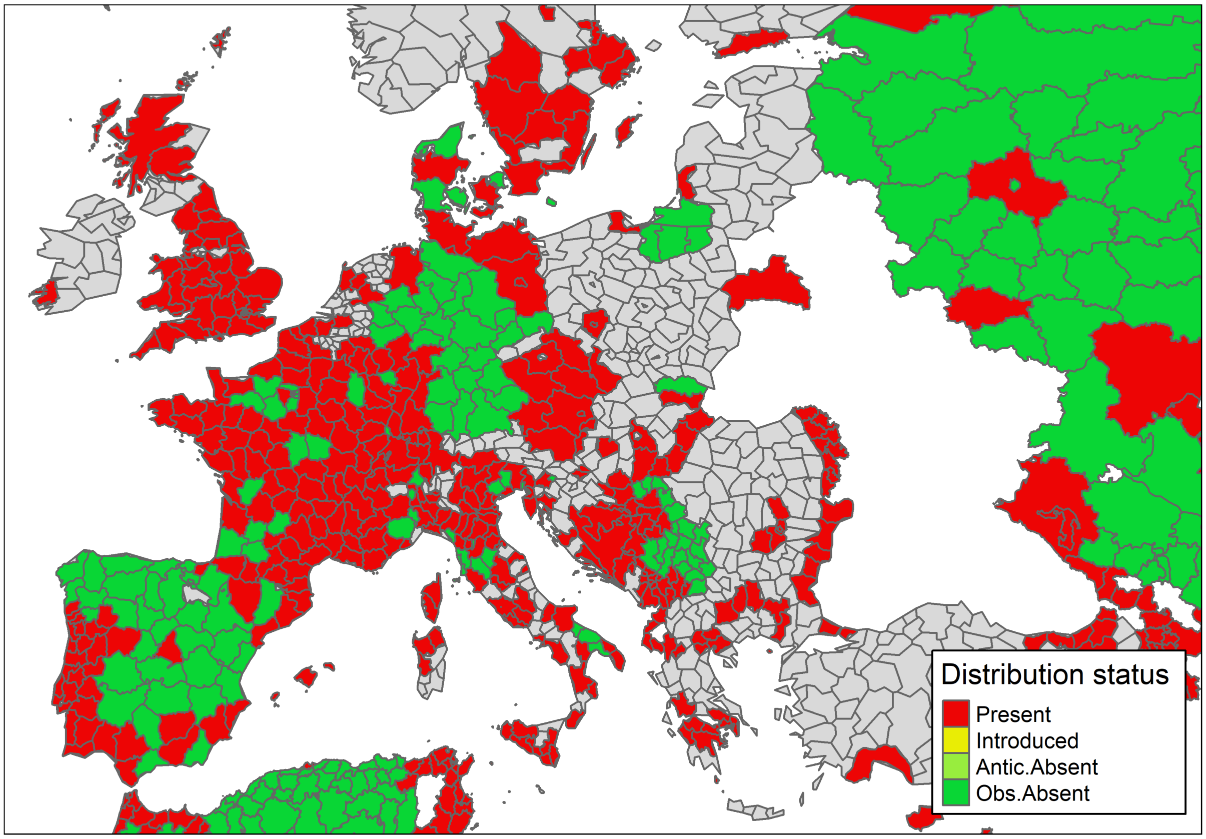


Figure S3: Confirmed Culex_pipiens NUTS3 distribution (source: ECDC).

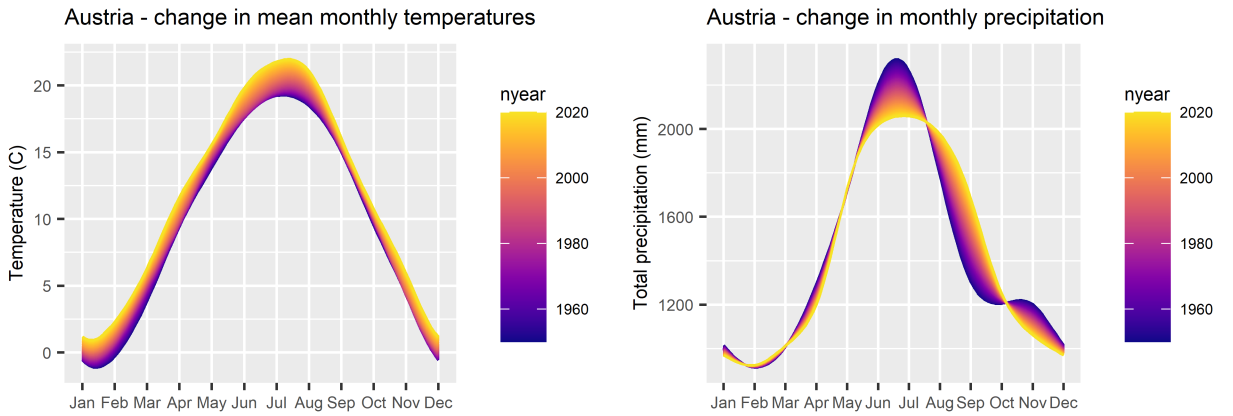


Figure S4: Austria - seasonal climate trends (Data source: E-OBS version 22.0e).

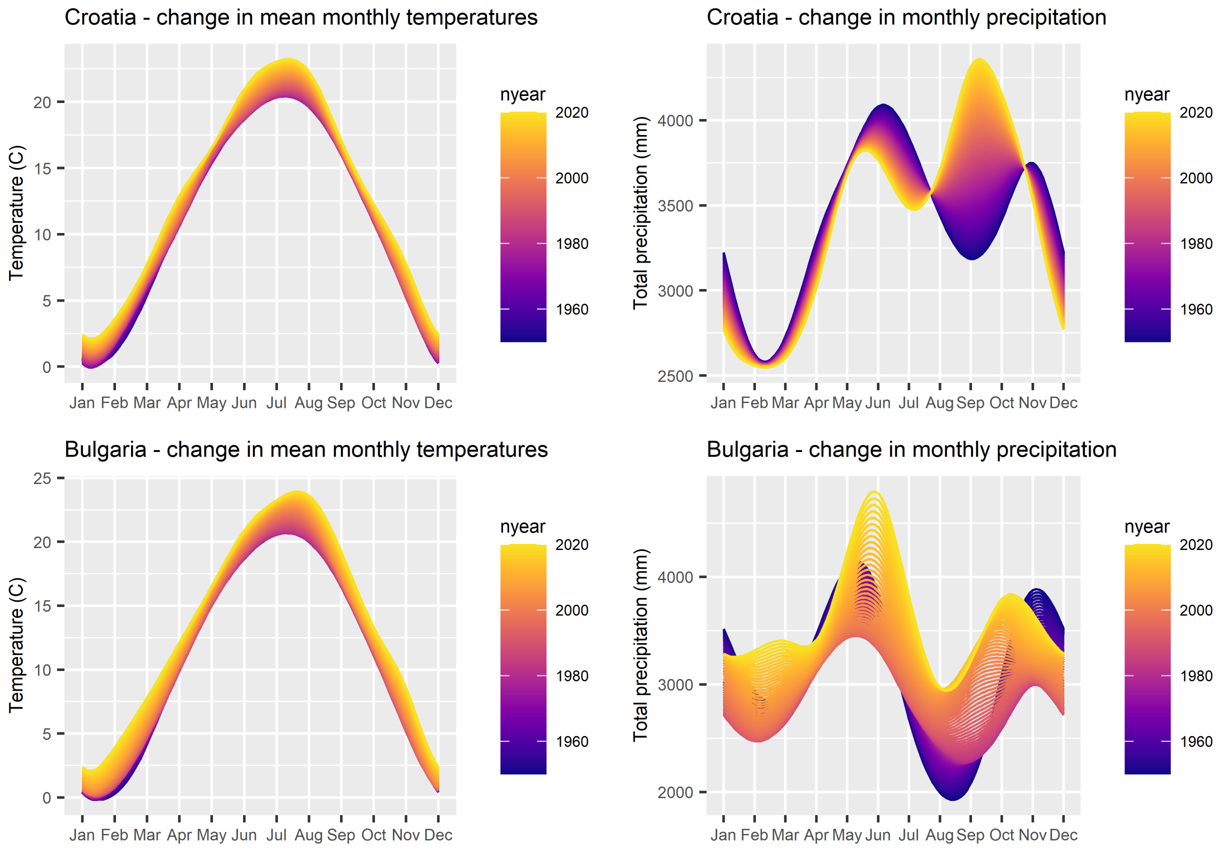


Figure S5: Bulgaria / Croatia - seasonal climate trends (Data source: E-OBS version 22.0e).

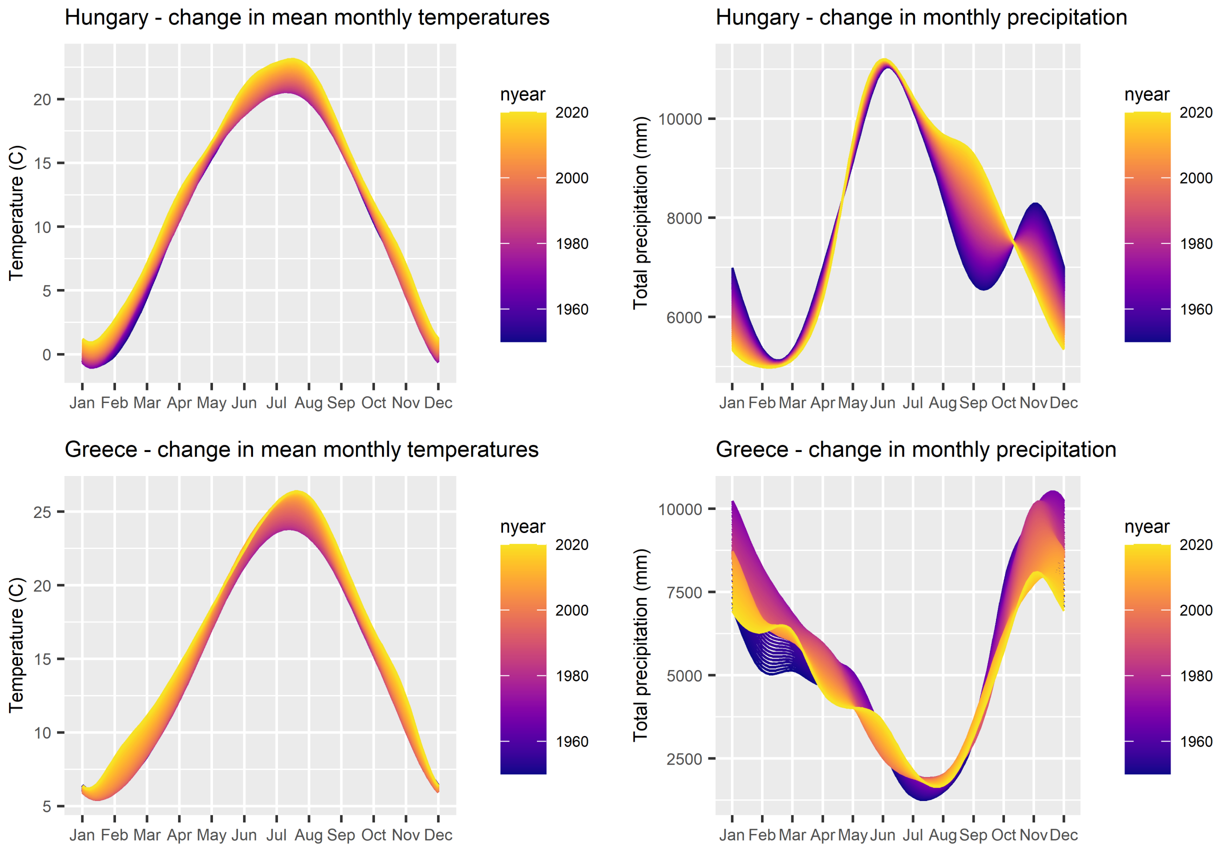


Figure S6: Greece / Hungary- seasonal climate trends (Data source: E-OBS version 22.0e).

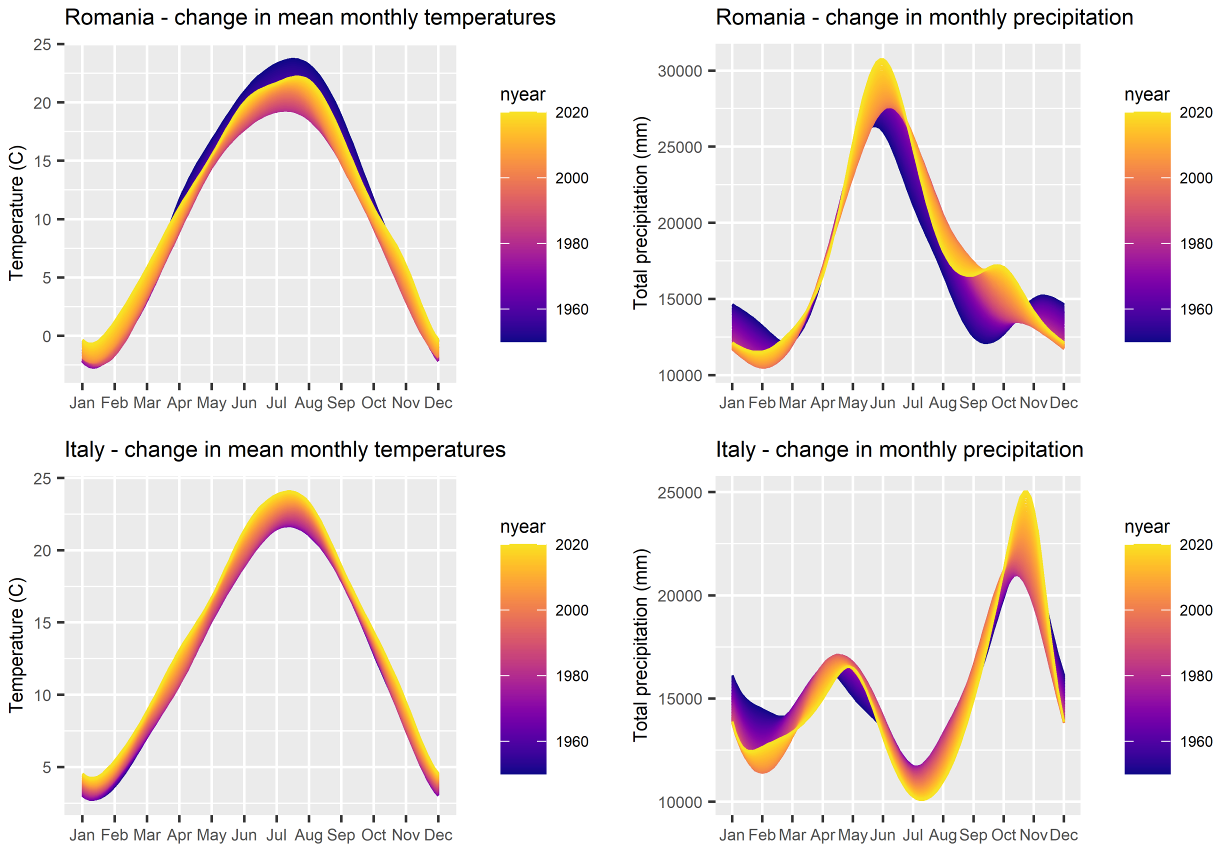


Figure S7: Romania / Italy - seasonal climate trends (Data source: E-OBS version 22.0e).

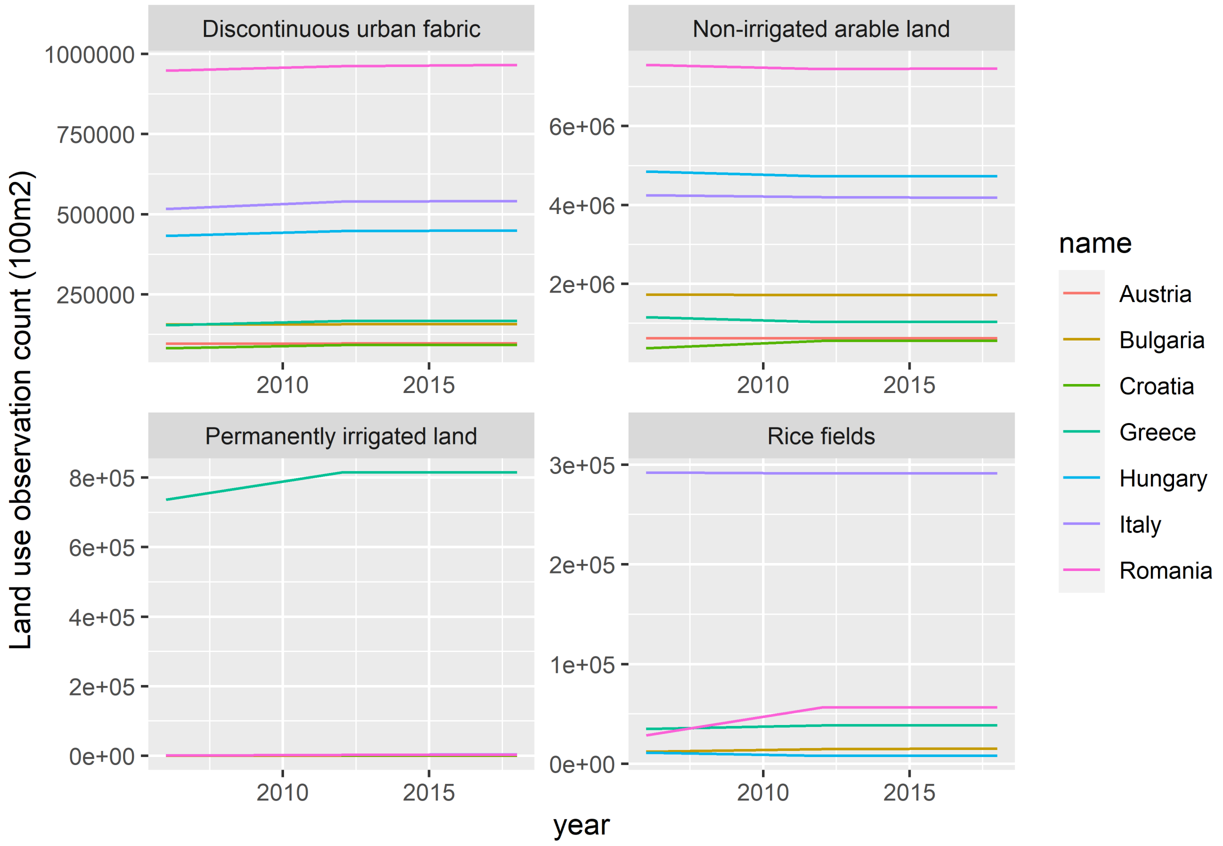


Figure S8: Land-use: 1 = Discontinuous Urban Fabric, 2-4 = Arable land break-down (Source: CORINE Land Cover)

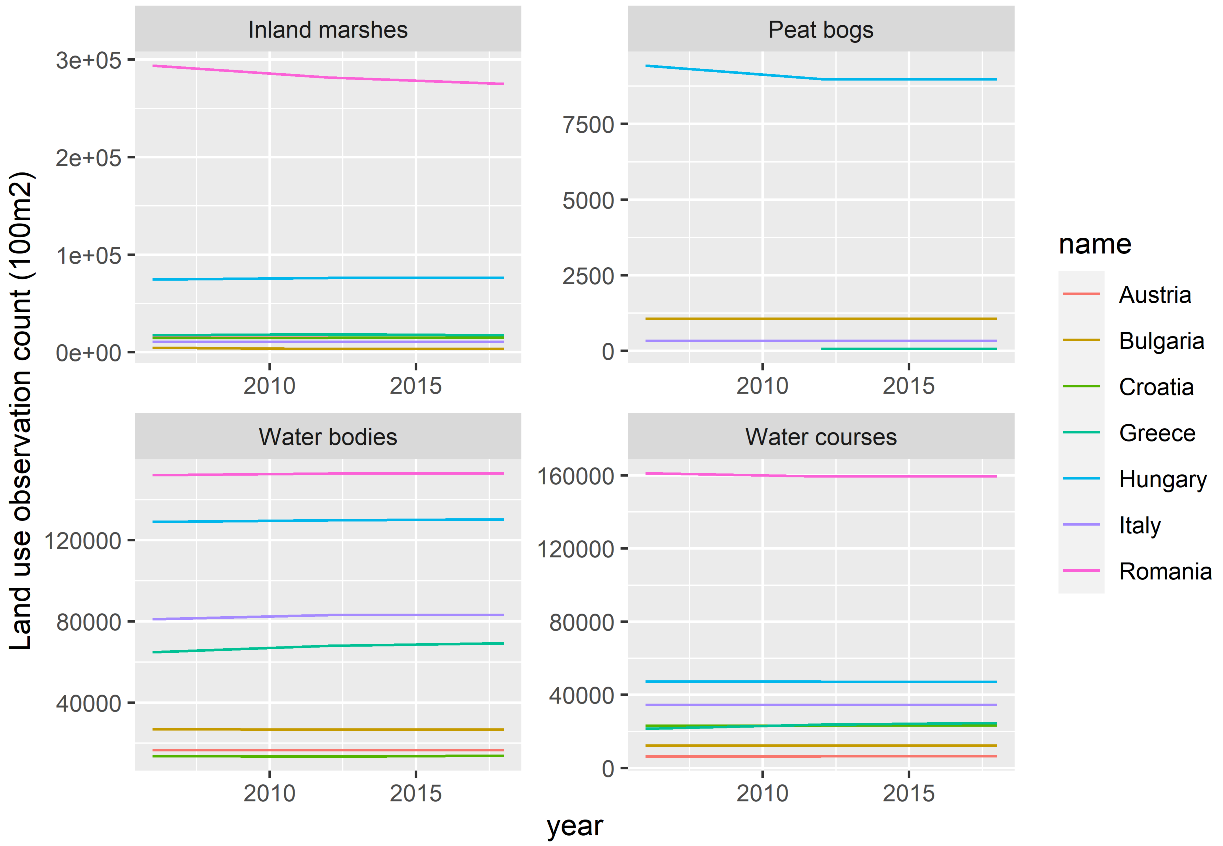


Figure S9: Land-use: Fresh water bodies break-down (Source: CORINE Land Cover)

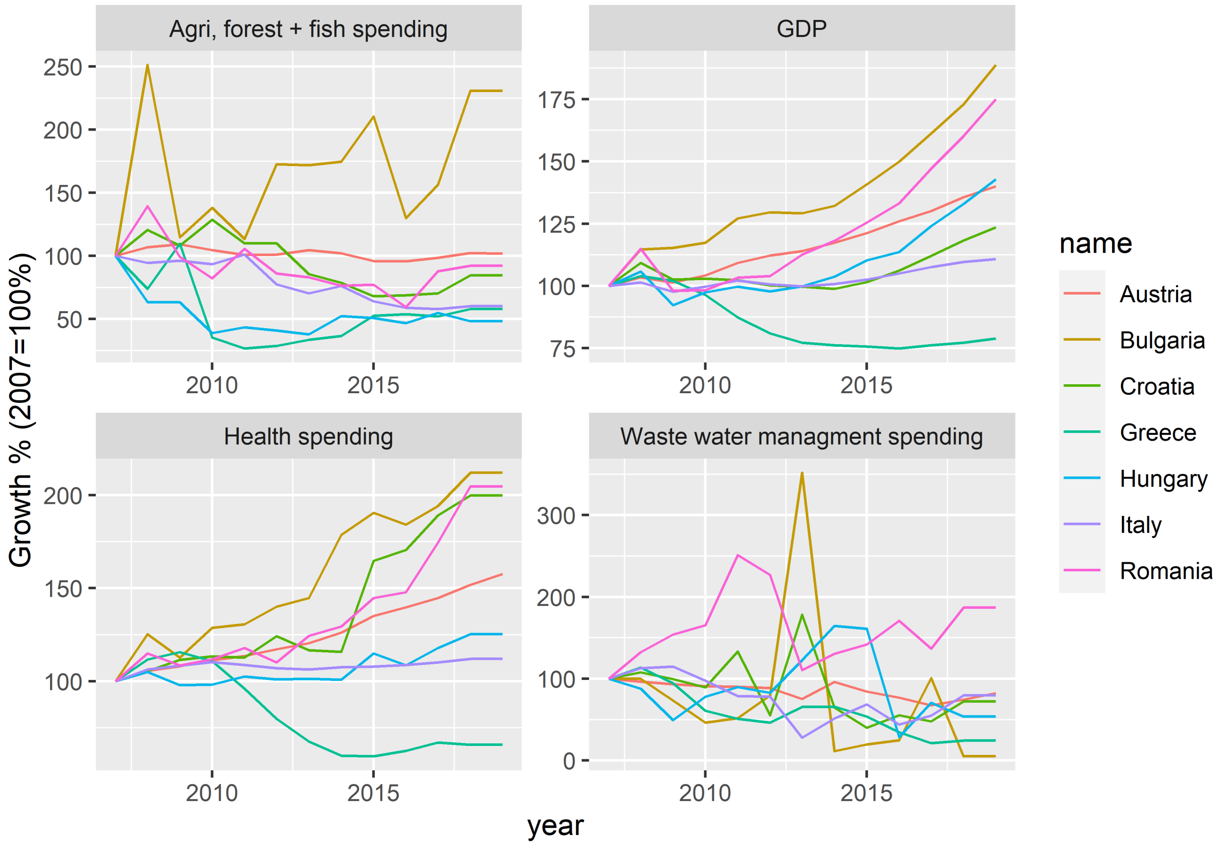


Figure S10: Government spending growth, GDP growth and unemployment 2007-2019 (2007=100%) (Source: Eurostat)

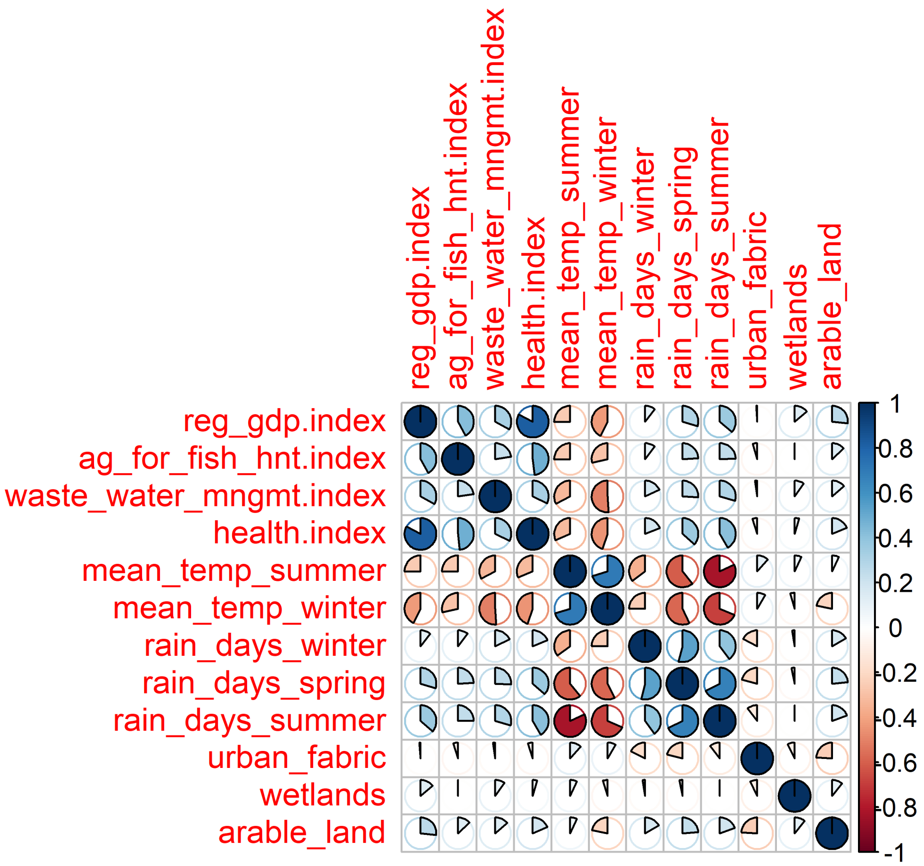


Figure S11: Variable correlation plot.

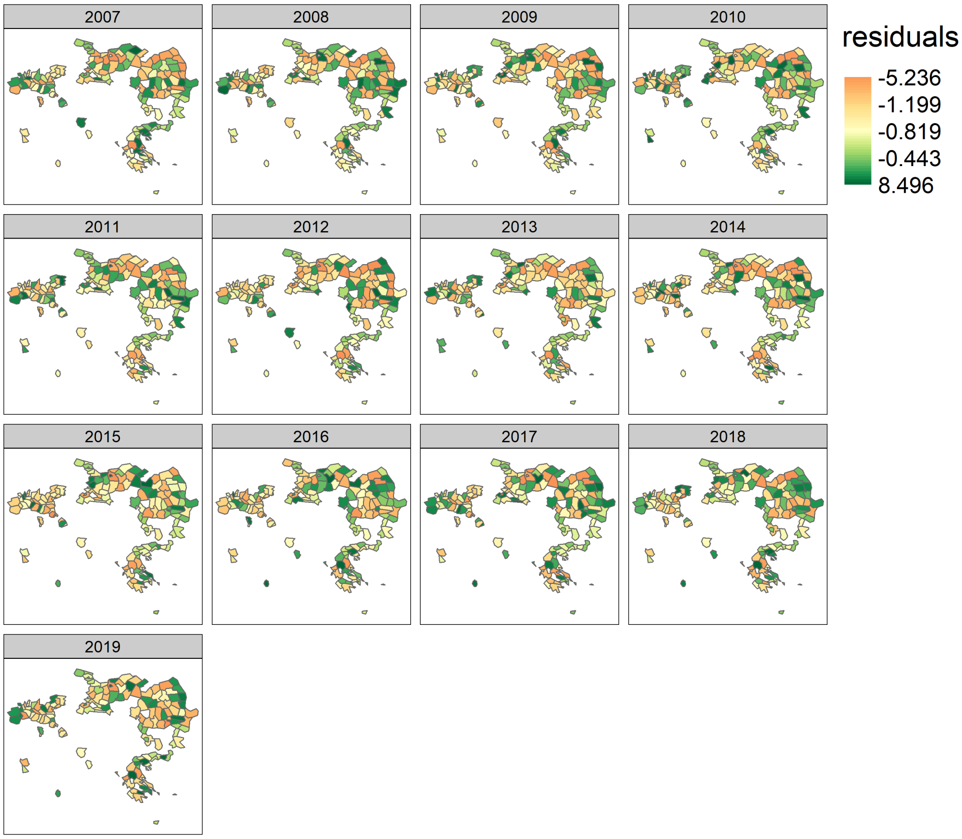


Figure S12: Spatial Residuals Tweedie model.

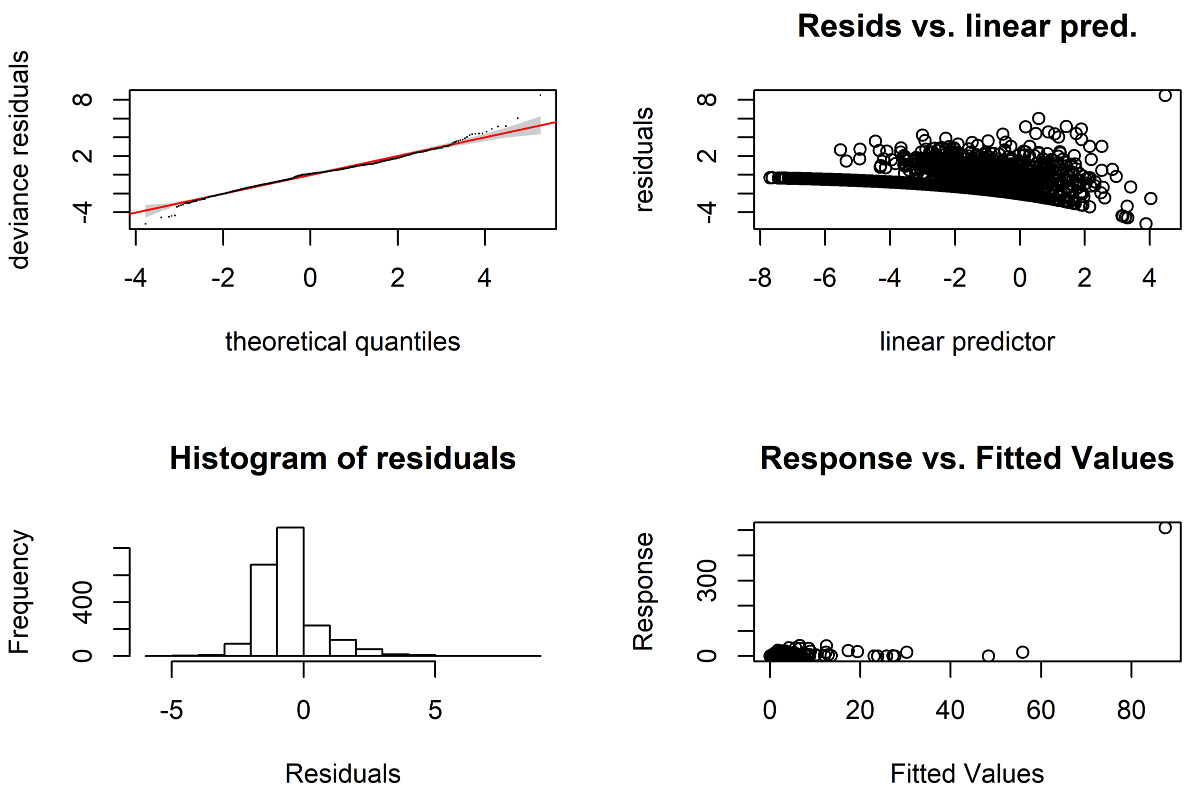


Figure S13: Diagnostics Tweedie model.

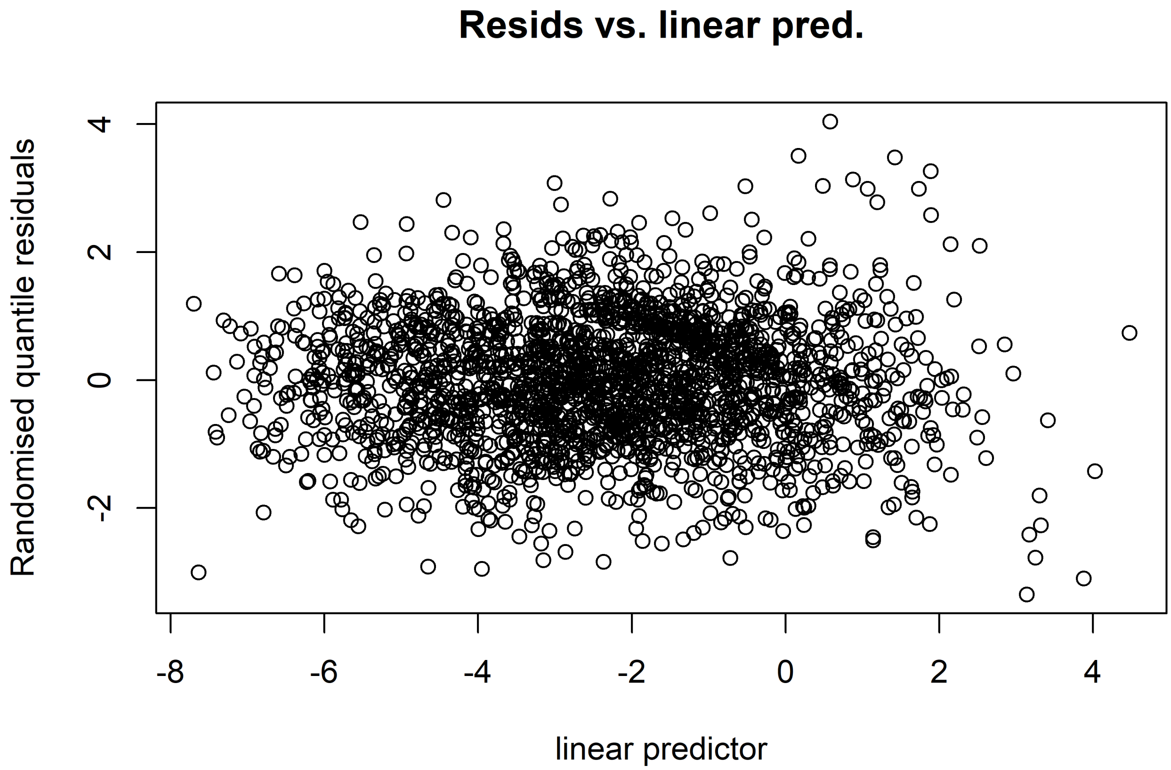


Figure S14: Diagnostics 2 Tweedie model.

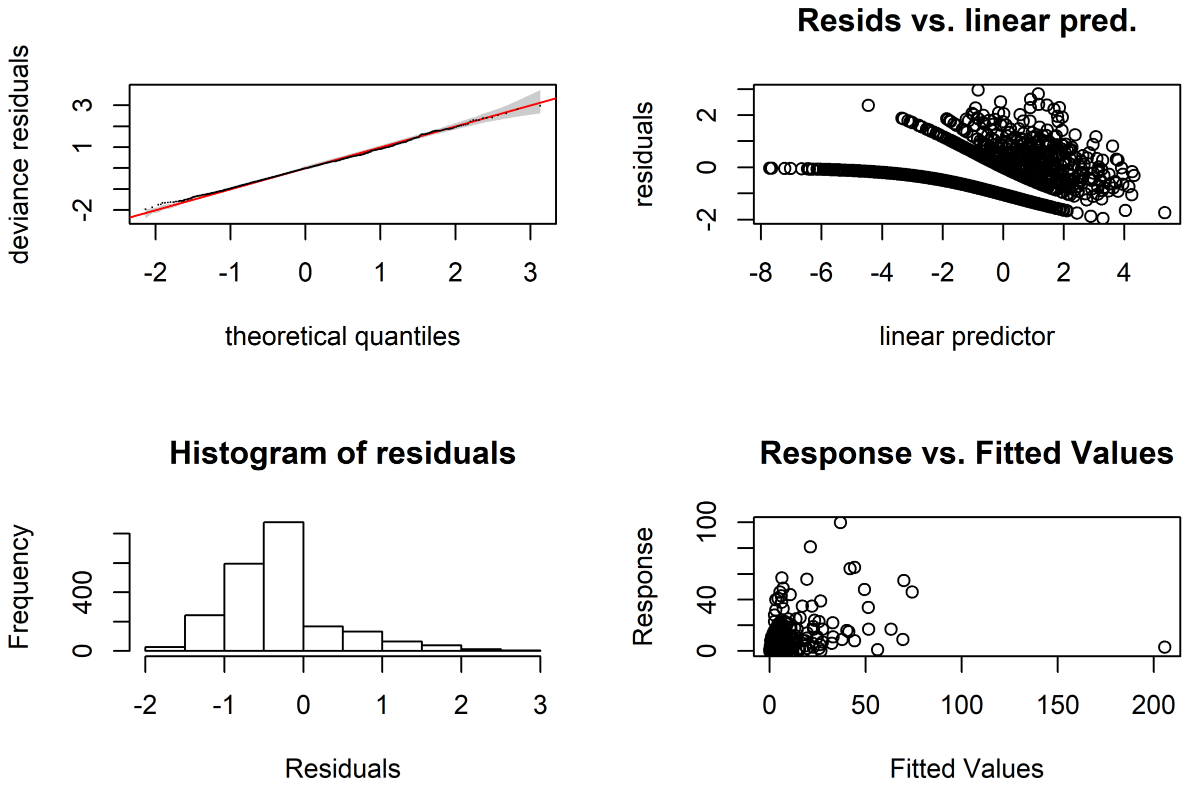


Figure S15: Diagnostics negbin model.

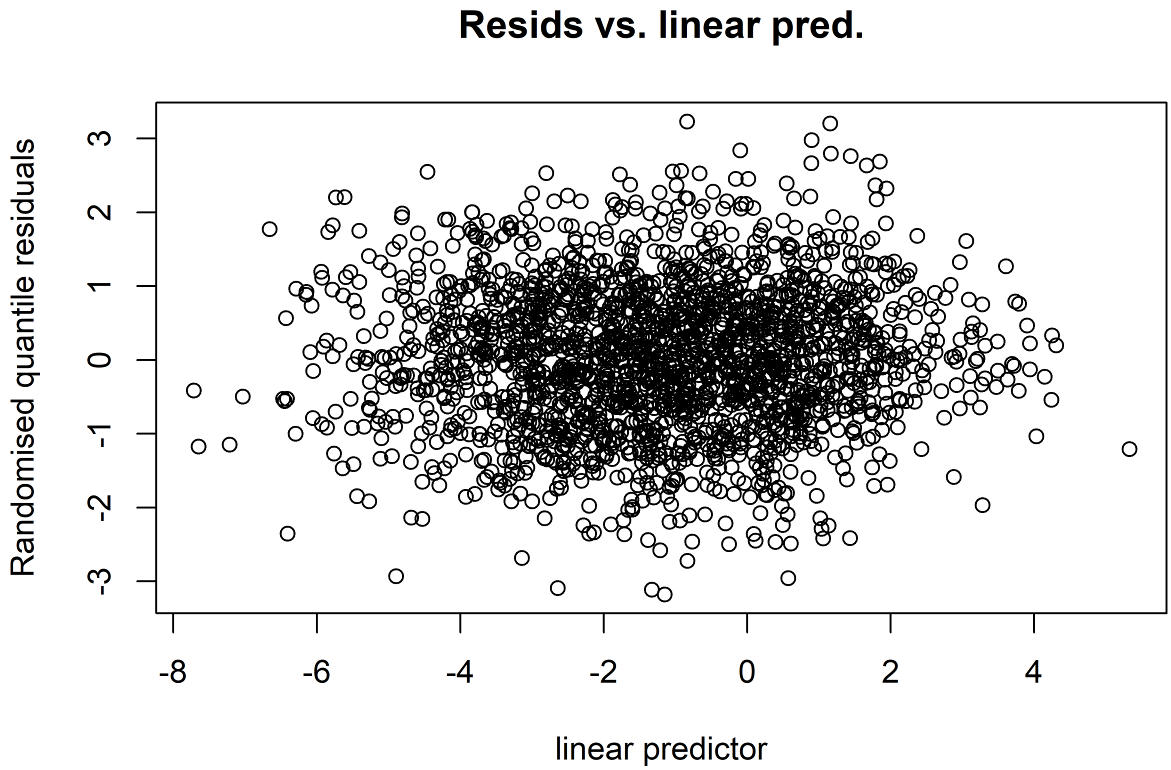


Figure S16: Diagnostics 2 negbin model.

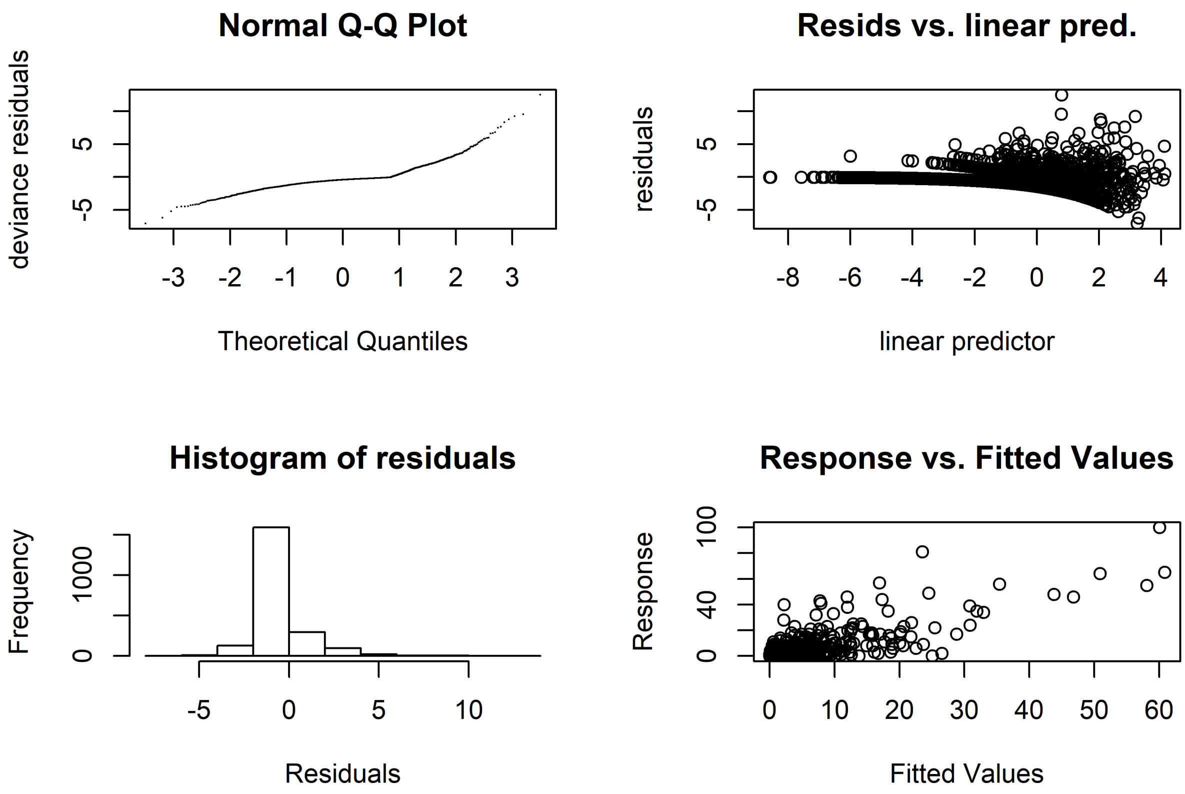


Figure S17: Diagnostics Quasipoisson model.

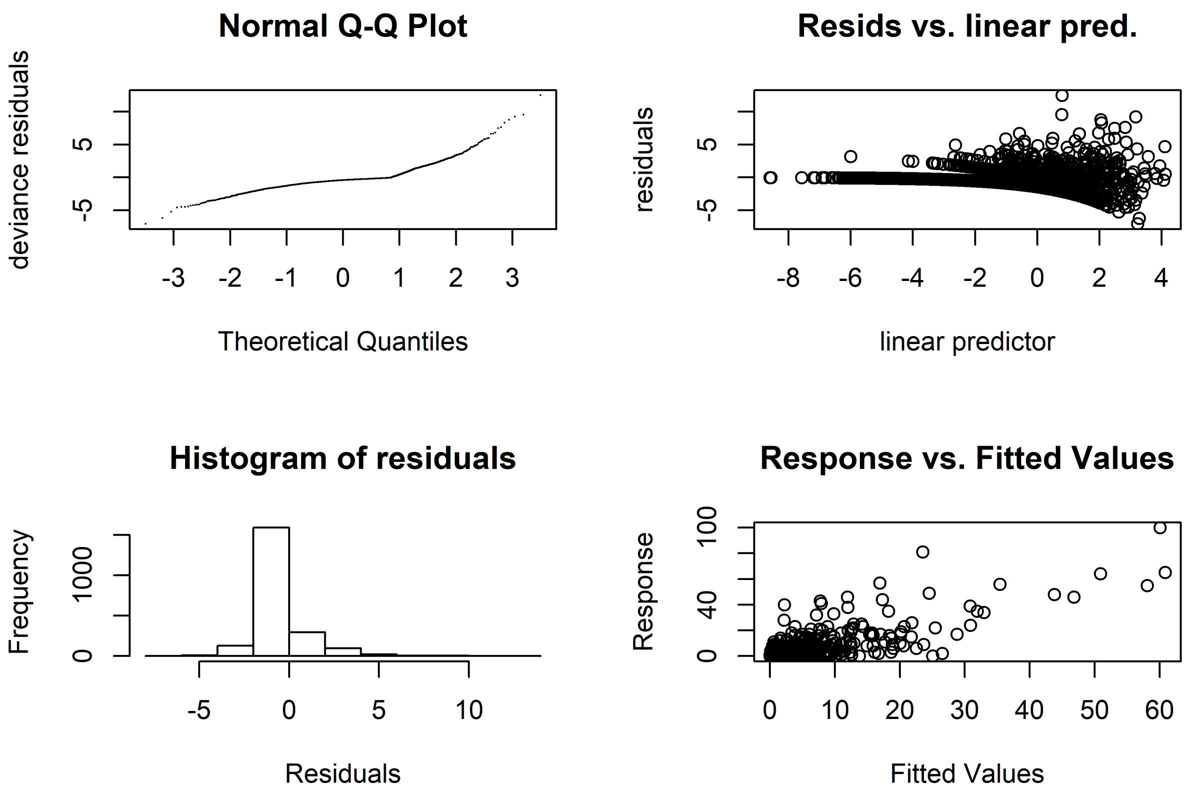


Figure S18: Diagnostics 2 Negbin model.

Table S1: WNF Cases Per Country 2006-2019

| Country | 2007 | 2008 | 2009 | 2010 | 2011 | 2012 | 2013 | 2014 | 2015 | 2016 | 2017 | 2018 | 2019 |
|---|---|---|---|---|---|---|---|---|---|---|---|---|---|
| Austria | 0 | 0 | 2 | 1 | 0 | 0 | 0 | 2 | 6 | 5 | 6 | 20 | 4 |
| Bulgaria | 0 | 0 | 0 | 0 | 0 | 2 | 0 | 1 | 2 | 1 | 1 | 15 | 6 |
| Croatia | 0 | 0 | 0 | 0 | 0 | 6 | 20 | 1 | 1 | 2 | 5 | 57 | 1 |
| Cyprus | 0 | 0 | 0 | 0 | 0 | 0 | 0 | 0 | 0 | 1 | 0 | 1 | 24 |
| Czechia | 0 | 0 | 0 | 0 | 0 | 0 | 1 | 0 | 0 | 0 | 0 | 5 | 1 |
| France | 0 | 0 | 0 | 0 | 0 | 0 | 0 | 0 | 1 | 0 | 2 | 27 | 1 |
| Germany | 0 | 0 | 0 | 0 | 0 | 0 | 0 | 0 | 0 | 0 | 0 | 1 | 4 |
| Greece | 0 | 0 | 0 | 262 | 100 | 157 | 85 | 15 | 0 | 0 | 48 | 312 | 228 |
| Hungary | 4 | 19 | 7 | 18 | 4 | 17 | 35 | 10 | 18 | 44 | 20 | 216 | 72 |
| Italy | 0 | 0 | 0 | 4 | 18 | 45 | 80 | 24 | 61 | 76 | 53 | 610 | 54 |
| Portugal | 0 | 0 | 0 | 0 | 0 | 0 | 0 | 0 | 1 | 0 | 0 | 0 | 0 |
| Romania | 4 | 2 | 2 | 57 | 11 | 15 | 24 | 23 | 32 | 93 | 66 | 279 | 68 |
| Slovakia | 0 | 0 | 0 | 0 | 0 | 0 | 0 | 0 | 0 | 0 | 0 | 0 | 1 |
| Slovenia | 0 | 0 | 0 | 0 | 0 | 0 | 1 | 0 | 0 | 0 | 0 | 4 | 0 |
| Spain | 0 | 0 | 0 | 2 | 0 | 0 | 0 | 0 | 0 | 4 | 0 | 0 | 0 |
| Turkey | 0 | 0 | 0 | 47 | 5 | 0 | 0 | 0 | 0 | 1 | 7 | 26 | 10 |

Table S2: Tweedie, negbin  Quasi Poisson model comparisons

| | Tweedie model | Negbin model | Quasi Poisson model |
|---|---|---|---|
| Intercept | $-2.35^{***}$ | $-13.82^{***}$ | $-13.86^{***}$ |
| | (0.40) | (0.36) | (0.45) |
| Mean temp summer (C) | $1.00^{*}$ | $1.50^{*}$ | $2.00^{**}$ |
| | (1.00) | (1.70) | (2.00) |
| Mean temp winter (C) | $1.94^{***}$ | $1.96^{***}$ | $2.00^{***}$ |
| | (1.99) | (1.99) | (2.00) |
| Days of rain in summer | $1.00^{**}$ | $1.00^{**}$ | 1.00 |
| | (1.00) | (1.00) | (1.00) |
| Summer surface water extent (30m2) | $1.02^{***}$ | $1.40^{*}$ | $1.62^{***}$ |
| | (1.03) | (1.64) | (1.85) |
| Continuous urban fabric % | $1.10^{***}$ | $1.45^{***}$ | $1.75^{***}$ |
| | (1.19) | (1.69) | (1.93) |
| Discontinuous_urban_fabric % | 1.00 | 1.00 | 1.00 |
| | (1.00) | (1.00) | (1.00) |
| Wetlands % | 1.00 | 1.00 | 1.00 |
| | (1.00) | (1.00) | (1.00) |
| Arable_land % | 1.00 | 1.00 | 1.01 |
| | (1.00) | (1.00) | (1.01) |
| Regional GDP index (2007=100%) | $1.74^{**}$ | $1.83^{***}$ | $1.00^{**}$ |
| | (1.84) | (1.90) | (1.00) |
| Agri, forest + fish spending (2007=100%) | $11.56^{***}$ | $11.55^{***}$ | $11.62^{***}$ |
| | (12.00) | (12.00) | (12.00) |
| Waste water managment spending (2007=100%) | $76.19^{***}$ | $83.79^{***}$ | $117.20^{***}$ |
| | (106.52) | (115.75) | (141.05) |
| Spatial lag | $1.93^{***}$ | $1.94^{***}$ | $2.00^{***}$ |
| | (1.99) | (1.99) | (2.00) |
| Year | 1.00 | 1.00 | 1.00 |
| | (1.00) | (1.00) | (1.00) |
| AIC | 3907.56 | 4659.28 | |
| BIC | 4538.85 | 5335.48 | |
| Log Likelihood | $-1842.57$ | $-2210.53$ | |
| Deviance | 3520.85 | 1207.27 | 4607.49 |
| Deviance explained | 0.65 | 0.64 | 0.69 |
| Dispersion | 2.73 | 1.00 | 3.22 |
| $R^2$ | 0.25 | $-0.32$ | 0.60 |
| GCV score | 1871.90 | 2367.22 | 2.45 |
| Num. obs. | 2158 | 2158 | 2158 |
| Num. smooth terms | 13 | 13 | 13 |

$^{***}p < 0.01$; $^{**}p < 0.05$; $^{*}p < 0.1$
